# An Atomistic Description of Heterotypic Lipid Exchange by Sec14-like Phosphatidylinositol Transfer Proteins

**DOI:** 10.64898/2026.07.20.739462

**Authors:** Tien Nguyen, Xiao-Ru Chen, Prashant Singh, Savana Green, M. Charles Cahill, Joy M. Shaffer, Danish Khan, Trivikram Molugu, Ashley Kidwell, Prasanna Iyer, Kiryl Zhaliazka, Sheena D’Arcy, Vytas A. Bankaitis, Tatyana I. Igumenova

## Abstract

Lipid transfer proteins (LTPs) are core regulators of the membrane dynamics, lipid signaling and intracellular communication networks that connect every organelle in the eukaryotic cell. ATP-independent lipid exchange reactions are a hallmark activity of these proteins. These remarkable reactions are essential for the important biological functions of LTPs but how lipid exchange is executed is not at all understood. Herein, we focus on phosphatidylinositol transfer proteins (PITPs) of the highly conserved and highly expanded Sec14/CRAL-Trio-like protein superfamily that potentiate phosphatidylinositol-4-phosphate (PtdIns4P) signaling in eukaryotic cells. Using an integrated structural approach, we describe in atomistic detail the lipid exchange reaction of Sec14-like PITPs. The molecular concepts we identify not only yield insights into how these PITPs integrate metabolic activity with PtdIns4P signaling in cells but also provide a framework for interpreting the functional mechanisms of other LTPs of the Sec14/CRAL-Trio superfamily.

## INTRODUCTION

Phosphatidylinositol (PtdIns) is the metabolic precursor for an essential class of phospholipids termed phosphoinositides (PIPs). These lipids are major intracellular signals in all eukaryotic cells. As such, their synthesis and degradation involves enzymes of critical biological importance.(Balla; Burke et al., 2023; Hammond and Burke, 2020; Posor et al., 2022; Swart and Hilbi, 2020; Vanhaesebroeck et al., 2021) Much effort is invested in deciphering the catalytic mechanisms of the highly conserved core enzymes that produce and consume PIPs as even subtle dysregulations of PIP metabolism are the root causes of multiple diseases in humans.(Pendaries et al., 2003; Volpatti et al., 2019) PtdIns transfer proteins (PITPs) garner intense interest from this perspective as these proteins impose a critical layer of additional regulation to PIP metabolism. PITPs do so by channeling PIP synthesis to specific biological outcomes and tuning PIP signaling to cellular metabolic needs.(Bankaitis et al., 2025; Bankaitis et al., 2010; Schaaf et al., 2008)

The fundamental biochemistry that underlies the PITP mode of action is rooted in the ability of these proteins to exchange a bound lipid for one in the cytosolic membrane leaflet at solution-membrane interfaces. PITPs do not fit the canonical definition of an enzyme because no chemical transformation of their lipid substrate(s) takes place during the exchange reaction. PITP-mediated lipid exchange reactions are also remarkable in their independence of energy input in vitro. How PITPs execute such complex and an ATP-independent interfacial processes remains unresolved. Herein, we combine X-ray crystallography, HDX-MS, solution NMR spectroscopy, atomistic MD simulations, and biochemical/genetic approaches to gain insight into the molecular mechanism of PITP function.

PITPs fall into two structurally unrelated families -- the Sec14/CRAL-Trio and the START -like PITPs.(Schaaf *et al*., 2008; Sha et al., 1998; Yoder et al., 2001) Sec14-like and START-like PITPs are expressed either as ‘soluble’ proteins or as membrane-tethering versions of inter-membrane contact sites.(Cockcroft and Lev, 2020; Kim et al., 2015; Kim et al., 2013; Milligan et al., 1997; Raghu et al., 2021) The soluble Sec14- and START-like PITPs execute important biological functions throughout the *Eukaryota*. These include essential roles in: (i) promoting membrane trafficking in yeast and mammalian neural stem cells, (ii) supporting critical developmental and stress response functions in plants, (iii) ensuring proper development of the mammalian forebrain and neural stem cell homeostasis, (iv) stimulating tumor growth/aggressiveness, and (v) preventing degenerative processes in mammalian central nervous and pancreatic insulin-secreting beta cell systems.(Bankaitis et al., 1990; Capitano et al., 2018; Hamilton et al., 1997; Hertle et al., 2020; Koe et al., 2018; Montag et al., 2023; Pathak et al., 2024; Vincent et al., 2005; Xie and Bankaitis, 2022; Yeh et al., 2023; Zhao et al., 2017)

Herein, we focus on Sec14 and its highly homologous paralog Sfh1 as model systems for addressing key mechanistic aspects of how PITPs execute their biological functions. Sec14 is a prototype for the Sec14-like PITP family and CRAL/Trio superfamily and represents the major yeast PtdIns/phosphatidylcholine (PtdCho) transfer protein. Sec14 activity is essential for yeast viability as it coordinates lipid metabolism and lipid signaling with biogenesis of protein transport vesicles on Trans Golgi Network/endosomal membranes and progression through the G1 phase of the cell cycle.(Bankaitis et al., 1989; Cleves et al., 1991; Huang et al., 2018; Mousley et al.) The principal basis of this cellular function rests on Sec14 potentiating PtdIns-4-phosphate (PtdIns4P) signaling by stimulating PtdIns 4-OH kinase activities.(Guo et al., 1999; Hama et al., 1999; Rivas et al., 1999)

Sec14 has a spacious lipid-binding cavity that harbors either PtdCho or PtdIns. As illustrated in **Fig. 1A**, upon membrane encounter Sec14 exchanges its resident lipid molecule for one it extracts from the bilayer. Whereas Sec14 executes both homotypic (PtdIns/PtdIns or PtdCho/PtdCho) and heterotypic (PtdCho/PtdIns or PtdIns/PtdCho) exchange reactions (**Fig. 1A**), it is during the course of the latter that Sec14 enhances PtdIns 4-OH kinase activity and executes its essential in vivo function.(Bankaitis *et al*., 2025; Bankaitis *et al*., 2010; Schaaf *et al*., 2008) Yet, the unique protein-lipid dynamics associated with lipid exchange reactions are not at all understood for Sec14, for other PITPs, or for LTPs in general. Herein, we provide an atomistic description for how Sec14-like PITPs execute heterotypic lipid exchange. We describe key features of this remarkable reaction and provide new insights into how Sec14-like PITPs regulate PtdIns4P signaling in cells. Finally, this report offers a high-resolution framework for conceptualizing lipid exchange mechanisms for other lipid transfer proteins.

**Figure 1.**
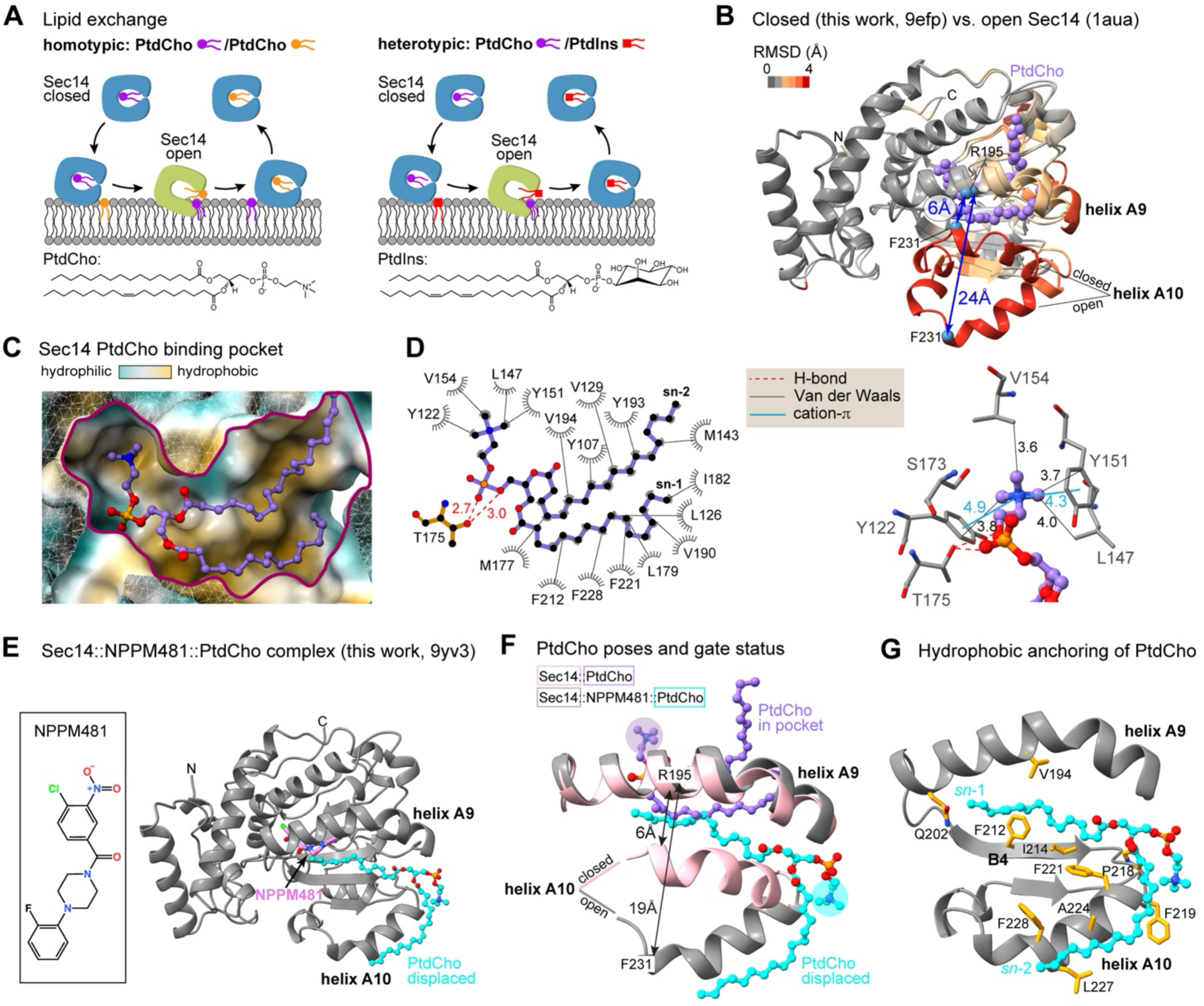
Open and closed conformations of Sec14. **(A)** Schematic view of Sec14 homotypic (PtdCho for PtdCho; left panel) and heterotypic (PtdIns for PtdCho; right panel) lipid exchange reactions illustrating conformational transitions between closed and open conformers. **(B)** Backbone superposition of the Sec14 open conformer (PDB 1AUA) and closed Sec14::PtdCho complex (PDB 9EFP, this work). R.m.s.d. values are color-coded and mapped onto both structures to illustrate gate opening. The gate ruler is defined as the distance between Cα atoms of R_195_ and F_231_. **(C)** A cut-away view of the bound PtdCho pose in the binding pocket of the Sec14::PtdCho crystal structure (PDB 9EFP, this work). PtdCho is shown in purple ball-and-stick representation. The pocket surface is color-coded according to hydrophobicity. **(D)** Summary of Sec14-PtdCho interactions, including van der Waals, hydrogen-bond, and cation–π contacts. **(E)** Structure of the Sec14::NPPM481::PtdCho ternary complex (PDB 9YV3, this work) with inhibitor NPPM481 and PtdCho shown in Van der Waals and ball- and-stick representations, respectively. **(F)** Comparison of PtdCho binding poses in the closed binary Sec14::PtdCho complex (gate distance 6 Å) and in the open ternary Sec14::NPPM481::PtdCho complex (gate distance 19 Å). **(G)** The NPPM481-displaced PtdCho is anchored to the protein by van der Waals interactions of its acyl chains with Sec14 residues of helix A10 and β-strand B4. The headgroup phosphate is fully solvent exposed.

## RESULTS

### Structural evidence for conformational plasticity of the Sec14 gate region during lipid exchange

Execution of the lipid exchange reaction requires the opening of the Sec14 lipid binding pocket to allow for lipid egress and uptake. To interrogate this complex reaction in more detail, we leveraged our discovery that we could use a potent small molecule inhibitor of Sec14 to trap a lipid exchange intermediate (i.e. Sec14::inhibitor::PtdCho ternary complex).(Chen et al., 2023) This capability made possible an unprecedented experimental examination of the protein and lipid transitions involved in this reaction by comparing the structure of an open Sec14 conformer(Sha *et al*., 1998) with those of the closed Sec14::PtdCho conformer and of a Sec14::inhibitor::PtdCho ternary complex.

Populating Sec14 during purification with its native lipid ligand PtdCho (absent in the heterologous expression host *E. coli*) enabled determination of a high-resolution structure of the ‘closed’ Sec14::PtdCho complex (PDB ID 9efp, **Table S1**). Comparison of the ‘closed’ Sec14::PtdCho structure with that of an ‘open’ Sec14 conformer identifies helices A9 and A10 as the major structural elements that control access to the lipid-binding cavity (**Fig. 1B**). In the closed conformer, A9 and A10 form an extensive hydrophobic interface (**Fig. S1A**). Using the Cα-carbons of A10 residue F_231_ and of A9 residue R_195_ as a distance ruler, that transition increases the F_231_/R_195_ distance from 6 Å in the closed conformer to 24 Å in the open (**Fig. 1B**). Disruption of the A9/A10 helical interface in the open conformer and the resultant 18 Å displacement of A10 allows access to the lipid binding cavity. These two Sec14 structures provide experimental support for prior inferences regarding an essential role of the A10 helix in lipid exchange.(Ryan et al., 2007; Schaaf *et al*., 2008; Sha *et al*., 1998)

In the closed Sec14::PtdCho complex, the lipid ligand takes full advantage of the amphiphilic nature of the Sec14 lipid binding cavity (**Fig. 1C**). PtdCho acyl chain methyls/methylenes and the choline headgroup methyls engage in van der Waals interactions with aliphatic sidechains that line the binding cavity. Polar interactions include H-bonds between the T_175_ hydroxyl and the headgroup phosphate oxygens, and cation-π interactions between the choline quaternary nitrogen and a Y_122_/Y_151_ sandwich (**Fig. 1D**). This arrangement recapitulates the structural features of the Sec14 paralog Sfh1 bound to PtdCho and of Sfh1^E126A^ bound to phosphatidylethanolamine (**Fig. S1B**).

The extensive networks of protein-lipid interaction and A9/A10 contacts suggest gate opening and PtdCho exit require significant rearrangements of the Sec14 elements that surround the lipid binding cavity. To examine this idea further, we took advantage of a potent small molecule inhibitor of Sec14 lipid-exchange activity (NPPM481) that displaces PtdCho from its Sec14 binding site when applied in a membrane-free system as demonstrated by solution NMR spectroscopy.(Chen *et al*., 2023) We used NPPM481 as a tool compound to prepare and crystallize a ternary Sec14::NPPM481::PtdCho complex that fulfils the operational criteria for a lipid exchange intermediate as it completes an NPPM481/PtdCho exchange reaction when presented with membrane.(Chen *et al*., 2023)

The structure of the ternary Sec14::NPPM481::PtdCho complex (PDB ID 9yv3; **Table S1**, **Fig. 1E**) differs from that of Sec14::PtdCho in key respects. First, the complex exhibits a partially open gate characterized by a ca.13 Å displacement of A10 relative to the closed conformer (**Fig. 1F**). Second, NPPM481 occupies the PtdCho headgroup binding region and displaces the lipid from its resident pose. The displaced PtdCho no longer engages in any of the protein interactions observed in the Sec14::PtdCho structure. Instead, its acyl chains make van der Waals contacts with helix A9 (V_194_, Q_202_), helix A10 (F_219_, F_221_, A_224_, L_227_, and F_228_) and β-strand B4 (I_214_, F_212_) of the lipid binding pocket floor (**Fig. 1G**). Since many of these interacting residues form the inter-helical A9/A10 interface in the closed Sec14 conformer (V_194_, F_221_, A_224_, L_227_, and F_228_; **Fig. S1A**), the structure suggests it is the acyl tails of the displaced PtdCho that induce and/or stabilize gate opening. The structure further indicates it is the solvent exposure of the displaced PtdCho that provides a thermodynamic incentive for the ternary complex to complete exchange of PtdCho for NPPM481 upon encountering membrane (**Fig. 1G**).

### ‘Closed’-gate PITP conformers are dominant in solution

Since gate motions might potentiate membrane association by Sec14/Sec14-like PITPs, we sought to determine if the gate region underwent conformational fluctuations in solution that result in exposure of lipid-binding regions. To that end, we carried out hydrogen/deuterium exchange (HDX)(Englander, 2006) experiments on the PtdCho complexes of Sec14 and of its paralog Sfh1, whose lipid exchange activity is significanlty reduced compared to Sec14.

Similar deuteration rates were observed for both proteins and their deuteration profiles were consistent with their structures as surface loops and helices exchanged more rapidly than buried β-strands (**Fig. S2A**). The lipid binding pockets of both Sec14::PtdCho and Sfh1:PtdCho showed very little to no deuteration, even after 10^5^ s of exchange. These regions include the β-strands that form the floor of the lipid-binding pocket and short regions of helices A8 and A9 proximal to the lipid headgroup (**Fig. S2B**). Although the moderate sequence identity between Sec14 and Sfh1 (64%) precludes quantitative comparisons, no obvious difference was detected in the deuteration profiles of the Sec14::PtdCho and Sfh1::PtdCho helical gates.

The HDX profiles of PtdCho complexes of Sfh1 and the ‘gain-of-function’ mutant Sfh1^E126A^ that has Sec14-like lipid exchange activities were also compared.(Schaaf et al., 2011) The E_126_A substitution causes local destabilization of helices A5 and A6 as evidenced by peptides from these helices incorporating more deuterium in Sfh1^E126A^ relative to Sfh1 (**Fig. S3A**). This effect correlates with loss of a hydrogen bond between sidechains of E_126_ and Y_109_ located at the end of helix A_6_ (**Fig. S3B**).(Schaaf *et al*., 2011) The increased deuteration of Sfh1^E126A^ becomes less localized after prolonged HDX, suggesting that destabilization of A5 and A6 enhances conformational flexibility of the overall fold. Nonetheless, E_126_A does not affect deuteration of helix A10 nor does it affect deuteration of structural elements involved in lipid binding (**Fig. S3C**). These HDX data indicate there is no significant population of ‘open’-gate PITP::PtdCho conformers in solution irrespective of their lipid transfer activity. The data further suggest that PITPs encounter membranes in the closed conformation.

### HDX reports on the conserved mode of PITP-membrane interactions

The next step was to identify the PITP regions that engage with membranes during the exchange reaction. Isotropically tumbling bicelles were used as membrane mimics. Bicelles support the Sec14 exchange activity and are compatible with both HDX and high-resolution NMR spectroscopy.(Chen *et al*., 2023) HDX data were collected for each of the three PITP::PtdCho complexes incubated with dimyristoylPtdCho (DMPC)/n-dodecyl-β-melibioside (DDMB) bicelles and each HDX profile was compared to the corresponding profile of that complex in solution (**Fig. 2A**).

**Figure 2.**
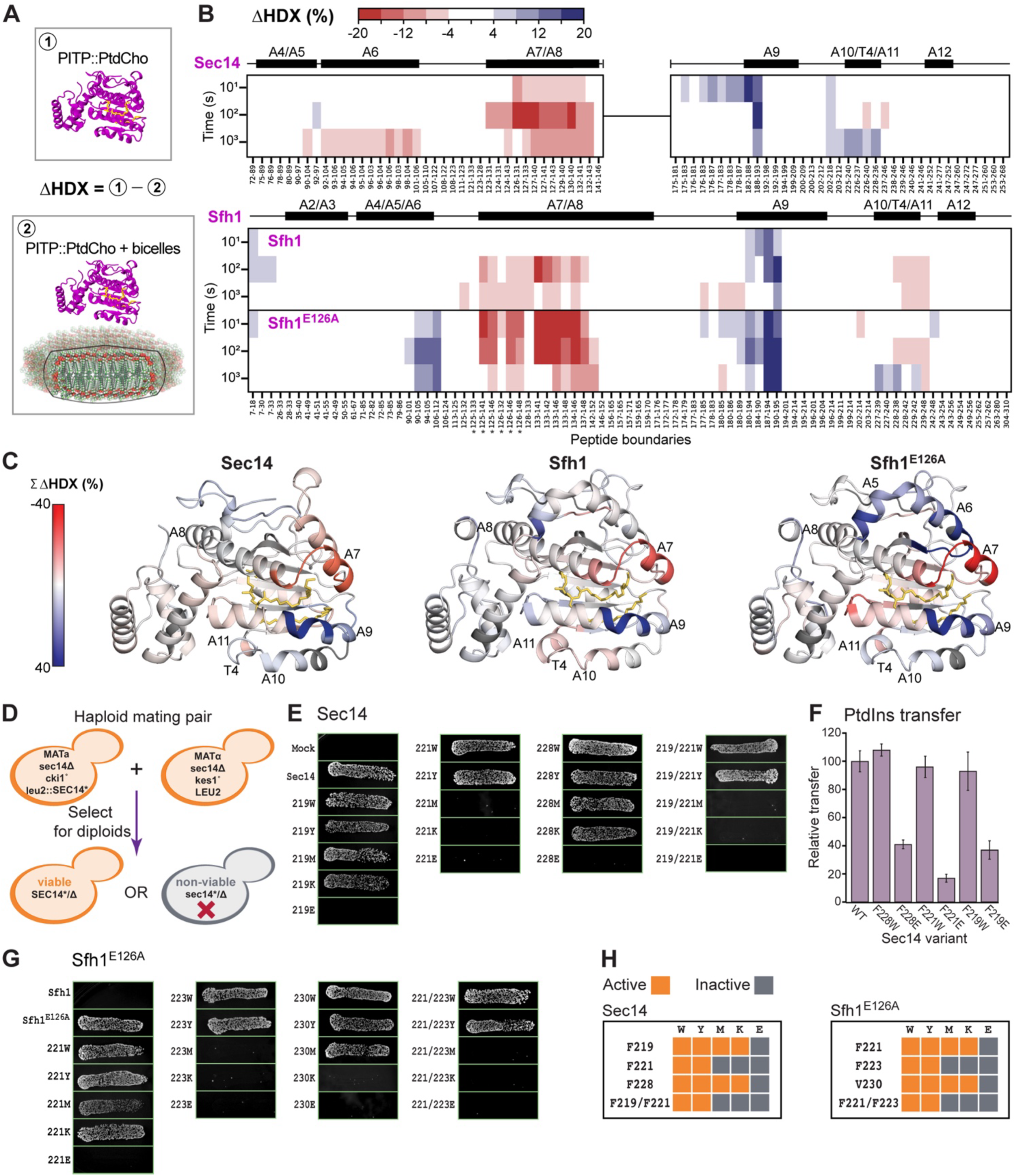
Putative membrane binding interfaces for Sec14::PtdCho, Sfh1::PtdCho, and Sfh1^E126A^::PtdCho. **(A)** Schematic representation of samples subjected to HDX measurements and definition of changes in deuteration upon bicelle addition (ΔHDX). **(B)** Time dependence of ΔHDX for Sec14::PtdCho, Sfh1::PtdCho, and Sfh1^E126A^::PtdCho with peptide identities shown on the x-axis. Peptides with ≥|4|% difference and a *p*-value ≤ 0.01 in a Welch’s t-test (n=2 or 3) are indicated in red (increase)/blue (decrease). Helices are numbered as in Sec14, and asterisks indicate peptides containing the E_126_A mutation. **(C)** Changes in deuteration upon the addition of bicelle mapped onto the structures of Sec14:: PtdCho (9EFP) or Sfh1:: PtdCho (3B7Q for Sfh1 and Sfh1^E126A^ data). Changes were summed across the 10, 10^2^, and 10^3^ s time points. Residue-level data were obtained from DynamX with no statistical filtering. Residues without coverage are shown in dark grey, and bound lipid is shown as yellow sticks. **(D)** Schematic representation of the ‘fatal attraction’ assays conducted on the PITPs and their variants (see Methods for details). Only a functional Sec14 variant supports diploid cell viability. **(E)** ‘Fatal attraction’ assay of Sec14 variants with substitutions of F_219_, and A10 helix residues F_221_ and F_223_. **(F)** The PtdIns-transfer activities of Sec14 F→W variants are comparable to that of the wild-type protein. Data represent the averages and standard deviations from 3 independent experiments. **(G)** ‘Fatal attraction’ assay of Sfh1^E126A^ variants with substitutions of F221, and A10 helix residues F_223_ and V_230_. Reconstitutions with Sfh1 and Sfh1^E126A^ expression serve as negative and positive controls, respectively. **(H)** Summary of biological activities of Sec14 and Sfh1^E126A^ variants in ‘fatal attraction’ assays.

The collective data report that Sec14, Sfh1, and Sfh1^E126A^ share a conserved mode of bicelle interactions as changes in deuteration (ΔHDX) occur in similar regions of these proteins (**Fig. 2B**). Bicelles reduce deuteration at the helix A9 N-terminal end and increase deuteration of helix A7 and part of helix A8. These profiles are consistent with A9 associating with the membrane surface and A7 engaging in enhanced solvent interactions as a result of membrane binding and/or lipid exchange of PtdCho for the shorter and more saturated DMPC. The kinetics of increased deuteration are similar for Sec14 and Sfh1^E126A^ but slower for Sfh1 – suggesting these differences reflect conformational transitions associated with lipid exchange.

The presence of bicelles also affected helix A10, with Sfh1 showing increased deuteration of the C-terminal end of A10, A11, and the loop that connects them (T4). Sec14 and Sfh1^E126A^ exhibit both increased and decreased deuteration of the A10/T4/A11 region depending on the time point (**Fig. 2C, Fig. S3D**). This hybrid effect suggests that some residues in the helix A10 peptides from Sec14 and Sfh1^E126A^ associate with the bilayer whereas others engage in enhanced solvent interactions. Of note, bicelle interaction overcomes the E_126_A-associated destabilization of Sfh1 helices A5 and A6 as evidenced by the corresponding reduction in deuteration and the similar rates with which bicelle-bound Sfh1^E126A^ and Sfh1 incorporated deuterium (**Fig. S3E**). Additional insight comes from a replicate HDX experiment with Sec14 that yielded improved coverage of helices A10/T4/A11 peptides (**Fig. S3F-H**). This experiment reports increased deuteration of T4 and A11, but reduced deuteration of the entirety of A10 and its preceding loop, in the presence of bicelles. In aggregate, the HDX data identify the N-terminal end of helix A9 and helix A10 as the Sec14/Sfh1 elements involved in membrane association.

### Gate mutations compromise Sec14-like PITP biological and biochemical activities

Sec14 helix A10 residues F_221_ and F_228_ are major contributors to formation of the A9/A10 hydrophobic interface in the closed Sec14 conformer (**Fig. S1A**). These residues, along with F_219_ of the A10 N-terminal loop, are also featured in PtdCho interactions within the ternary Sec14::NPPM481::PtdCho complex (**Fig. 1G**). Consequently, these residues represent attractive candidates for involvement in lipid exchange.

To address whether F_219_, F_221_, and F_228_ are important for Sec14 function in vivo, we devised a ‘fatal attraction’ mating assay whose readout for a functional Sec14 is recovery of viable diploid cells (**Fig. 2D**). As expected, viable diploids are recovered in positive control mating arrangements (functional Sec14 expression reconstituted) but not in mock control matings (no Sec14 activity expressed; **Fig. 2E)**. Test matings demonstrate F→E substitutions in helix A_10_ yield biologically non-functional Sec14 proteins. F_219_→K and F_228_→K are tolerated likely because of lysine’s ability to “snorkel” in the membrane environment. The F_221_ position is particularly sensitive to substitutions other than aromatic amino acids. PtdIns transfer assays with purified proteins provide biochemical confirmation that the biological insufficiencies of the Sec14 mutants are associated with defects in their lipid exchange activities (**Fig. 2F**).

The functional distinction between Sfh1 and Sfh1^E126A^ is on display in these ‘fatal attraction’ assays as Sfh1 expression fails to support diploid cell viability whereas Sfh1^E126A^ expression does so effectively (**Fig. 2G**). Moreover, the results of Sfh1^E126A^ mutagenesis experiments recapitulate those scored for Sec14 (**Fig. 2H**). These data: (i) suggest Sec14/Sfh1/Sfh1^E126A^ bind membranes similarly, in full agreement with the HDX results, and (ii) demonstrate that A10 plays an important role(s) in lipid exchange and in vivo function.

### ^19^F NMR reports on the formation of PITP-bicelle complexes

To directly probe PITP interactions with membranes, we devised a ^19^F-based solution NMR approach that (i) relies on the detection of PITP aromatic residues, and (ii) employs isotropically tumbling DMPC/dipalmitoyl-PtdCho (DHPC) bicelles as membrane mimics. ^19^F is a highly sensitive spin ½ nucleus whose chemical shift is exquisitely responsive to changes in its electronic environment.

^19^F labels were introduced into the three PITPs by metabolically substituting all native Trp residues with a fluorinated 5-^19^F-Trp analog. The fluorinated proteins were purified as Sfh1::PtdEtn(Schaaf et al., 2006), Sfh1^E126A^::PtdCho, and Sec14::PtdCho complexes (see Experimental Procedures), and their lipid transfer activities were not affected by fluorination (**Fig. S4A**). The NMR spectra show the expected number of spectrally resolved ^19^F signals -- 3 for Sec14 (W_82_, W_149_, and W_285_) and 2 for Sfh1/Sfh1^E126A^ (W_78_ and W_287_). Upon addition of bicelles, a uniform broadening of the ^19^F peaks for all three proteins was observed (**Fig. 3A**, blue spectra). This line broadening is caused by an increase of the rotational correlation time that directly reports on the formation of the PITP-bicelle complexes. Because native Trp residues are not part of the membrane-binding interface, their ^19^F chemical shift changes are minor and only affect the W_78_ site for Sfh1 and Sfh1^E126A^ (**Fig. 3A**). We attribute the 0.5 (0.4) ppm upfield shift of the Sfh1 (Sfh1^E126^) W_78_ ^19^F peak to the change in local environment caused by the exchange of resident lipid for one of the bicelle lipids (**Fig. S4B**).

**Figure 3.**
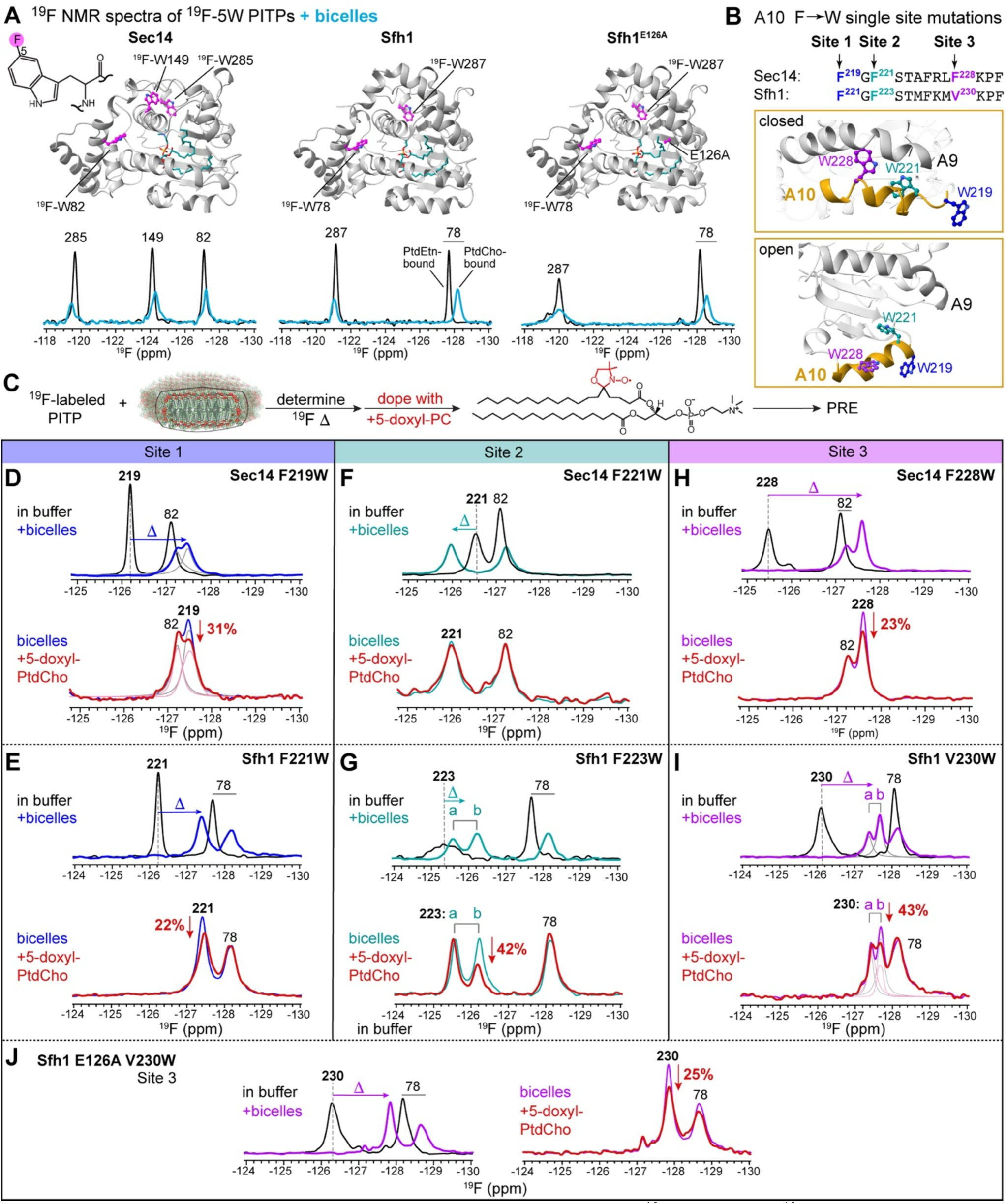
Sec14-like PITP membrane interactions probed by ^19^F NMR. **(A)** ^19^F NMR spectra of Sec14, Sfh1, and Sfh1^E126A^ metabolically labeled with 5-fluorotryptophan in the absence (black) and presence (blue) of DMPC/DHPC bicelles. Uniform peak broadening upon bicelle addition reports on the formation of protein-bicelle complexes. PtdCho from bicelles replaces co-purified PtdEtn. **(B)** Single-site Sec14-like PITP mutants chosen for incorporation of ¹⁹F-Trp probes: Site 1, F_219_^Sec14^/F_221_^Sfh1^; Site 2: F_221_^Sec14^/F_223_^Sfh1^; and Site 3: F_228_^Sec14^/V_230_^Sfh1^. The site positions are mapped onto helix A10 of Sec14 in closed and open conformations for visualization purposes,. **(C)** Experimental workflow for assessment of protein-membrane interactions and depth of insertion. Δ is the ^19^F site-specific chemical shift perturbation due to bicelle interactions. PRE is the % decrease of ^19^F peak intensity due to proximity to the membrane-embedded paramagnetic agent. **(D,E)** Δ and PRE data for Site 1 of Sec14 and Sfh1. Site 1 of both proteins inserts into the membrane as evidenced by their >20% PRE values. **(F,G)** Δ and PRE data for Site 2 of Sec14 and Sfh1. Site 2 in Sec14 interacts with membranes but peripherally as there is no detectable PRE (**F**). Site 2 of Sfh1 adopts two conformations (‘a’ and ‘b’) upon membrane binding. Only conformation ‘b’ inserts into the membrane (**G**, 42% PRE). **(H, I, J)** Δ and PRE data for Site 3 of Sec14, Sfh1, and Sfh1^E126A^. Site 3 undergoes membrane insertion in Sec14 (**H**, 23% PRE) and Sfh1^E126A^ (**J**, 25% PRE). Site 3 in Sfh1 shows similar spectroscopic signatures to those of Site 2 as it adopts conformations ‘a’ and ‘b’ upon bicelle interaction. Only conformation ‘b’ inserts into the membrane (**I**, 43% PRE). Spectra containing closely spaced peaks were deconvoluted using Line Fitting tool in MestReNova.

### A10 inserts into the bilayer and undergoes conformational exchange

To directly examine whether Sec14/Sfh1 helix A10 is a primary component of the PITP::membrane interaction interface, we introduced three single-site F→W mutations across helix A10 and labeled the proteins with 5-^19^F-Trp. For convenience, these mutant sites are referred to as Site 1 (F_219_W in Sec14, F_220_W in Sfh1), Site 2 (F_221_W in Sec14, F_223_W in Sfh1), and Site 3 (F_228_W in Sec14, V_230_W in Sfh1) (**Fig. 3B**). Biochemical assays demonstrate that all 5-^19^F-Trp-substituted Sec14 and Sfh1 proteins exhibit wild-type lipid transfer activities in vitro (**Fig. S4A**).

The PITP ^19^F NMR spectra were collected under three experimental conditions: (i) in buffer, (ii) in the presence of diamagnetic DMPC/DHPC bicelles, and (iii) in the presence of paramagnetic DMPC/DHPC bicelles doped with 5-doxyl-PtdCho (**Fig. 3C**). The NMR observables reporting on PITP-membrane interactions are ^19^F chemical shift perturbations (Δ) and paramagnetic relaxation enhancement (PRE). Δ values reflect changes in the ^19^F electronic environment due to interactions with bicelles and are obtained from comparative analysis of NMR spectra (i) and (ii). PRE values report on the depth of ^19^F insertion into the bilayer. Spatial proximity of the ^19^F nucleus to the 5-doxyl-PtdCho spin-label results in faster relaxation and hence decrease in NMR peak intensity. PRE is calculated from spectra (ii) and (iii) as the % decrease of the ^19^F peak intensity in paramagnetic relative to diamagnetic bicelles. Thus, ^19^F-labeled Trp sidechains of PITPs that insert into the lipid bilayer are expected to have non-zero values for both Δ and PRE.

Site 1 resides in the loop region that connects strand B4 with helix A10 and immediately precedes the beginning of A10. Both ^19^F-W_219_^Sec14^ and ^19^F-W_221_^Sfh1^ give rise to sharp peaks at −126.2 ppm in buffer indicating significant mobility of Trp sidechains at Site 1 -- as expected from their loop position (**Fig. 3D,E**). Upon addition of diamagnetic bicelles, the ^19^F-W_219_^Sec14^ and ^19^F-W_221_^Sfh1^ peaks broaden and shift upfield by 1.1 ppm and 1.2 ppm, respectively. In paramagnetic bicelles, Site 1 ^19^F peak intensities are attenuated by 31% for W_219_^Sec14^ and 22% for W_221_^Sfh1^. No such attenuation in intensity is observed for the ^19^F signals of native Trp residues (**Fig. 3D,E**). Together, large chemical shift changes Δ and >20% PRE values report that Site 1 inserts into the membrane.

Site 2 is located at the N-terminus of helix A10 and is a key component of the hydrophobic A9/A10 interface. The ^19^F-W_221_^Sec14^ peak shifts 0.55 ppm downfield in the presence of bicelles but shows no PRE (**Fig. 3F**). We conclude that Site 2 in Sec14 engages membrane primarily in peripheral interactions. The Sfh1 ^19^F-W_223_^Sfh1^ peak is significantly broadened in bicelle-free conditions (**Fig. 3G**). ^19^F Carr-Purcell-Meiboom-Grill (CPMG) relaxation dispersion experiments demonstrate this broadening is due to the μs-ms dynamics of the W_223_^Sfh1^ sidechain (**Fig. S4C**). Addition of bicelles results in the appearance of two peaks that we assign to different W_223_^Sfh1^ sidechain conformers designated as ‘a’ (−125.6 ppm) and ‘b’ (−126.2 ppm). Remarkably, only the upfield ‘b’ conformer shows strong PRE, while ‘a’ is unaffected by the presence of the paramagnetic lipid. Collectively, these data suggest that the A9/A10 helical interface accommodates sidechain dynamics, and that the aromatic sidechain of Site 2 in A10 can adopt both peripheral and inserted membrane geometries.

Site 3 is located in the middle of helix A10 and, similarly to Site 2, is a major player in the formation of the A9/A10 helical interface. Addition of bicelles results not only in a large upfield shift (Δ=2.1 ppm) of the ^19^F-W_228_^Sec14^ resonance (**Fig. 3H**), but also in the 1.5-fold reduction in its linewidth, as reported by the ^19^F transverse relaxation time constants T_2_ (**Fig. S4D**). This reduction in linewidth reports a significant increase in sidechain mobility upon interaction with bicelles. In addition to altered dynamics, Site 3 undergoes membrane insertion as evidenced by the 23% PRE of the ^19^F-W_228_ ^Sec14^ peak (**Fig. 3H**). In the case of Sfh1, Site 3 shows the same spectroscopic signatures as Site 2: (i) a broadening of the ^19^F-W_230_^Sfh1^ peak due to μs-ms dynamics in the bicelle-free system (**Fig. S4C**), (ii) an upfield shift upon interaction with bicelles; and (iii) appearance of two ^19^F-W_230_^Sfh1^ sidechain conformers ‘a’ and ‘b’, of which only the ‘b’ conformer inserts into the bilayer (**Fig. 3I**).

In aggregate, the ^19^F NMR data of **Figs. 3D-I** produce the following conclusions regarding the A10 helix of these PITPs. First, A10 inserts into the membrane, and the insertion event is accompanied by partial or full gate opening. Indeed, insertion of Site 2 and 3 aromatic sidechains into the bilayer is only possible upon disruption of the A9/A10 helical interface with resultant exposure of the lipid binding pocket (**Fig. 3B**). Second, membrane association alters A10 dynamics. The experimental manifestations of such are the increased mobility of Sec14 Site 3 (**Fig. 3H**) and the formation of two slowly inter-converting (see below) conformers of Sites 2 and 3 in Sfh1 (**Fig. 3G,I**). In the ‘gain-of-function’ Sfh1^E126A^ mutant, we were only able to probe Site 3 since 5-^19^F-Trp substitution of Trp residues at Sites 1 and 2 destabilizes these proteins when produced in *E. coli*. Remarkably, we find that the ^19^F-W_230_^Sfh1E126A^ NMR peak shows the same spectroscopic signatures as Sec14 Site 3: a large upfield chemical shift of 1.5 ppm upon addition of bicelles, a reduction in linewidth (**Fig. S4D**), and a 25% PRE (**Fig. 3J**). Since Sec14/Sfh1^E126A^ and Sfh1 are fast and slow lipid exchange PITPs, respectively, we hypothesized that A10 dynamics directly reports on the gate-opening motions that enable lipid exchange process in these Sec14-like PITPs.

### Membrane association is required for gate opening motions

To determine if NMR-detected A10 dynamics reports on the gate movement, we designed Sfh1 and Sec14 variants that ‘lock’ the proteins in a closed form. This was accomplished by engineering Cys substitutions into helices A9. and A10 and cross-linking the sites under non-reducing conditions. We successfully crystallized and determined a 1.8 Å structure of the Sfh1^F223W,K197C,F233C^ mutant (Sfh1^lock^, PDB 9efn). The structure is of a closed conformer with the gate covalently locked via the expected disulfide bond and a PtdEtn bound inside the lipid binding cavity (**Fig. 4A; Table S1**). No structural distortions due to disulfide bond formation were observed (backbone r.m.s.d. between the cross-linked structure and that of the Sfh1::PtdEtn complex (PDB 3b74) is 0.3 Å). While Sfh1^lock^ is inactive for PtdIns-transfer under non-reducing or mildly reducing conditions, its activity is fully restored by high concentration of reductant (**Fig. 4B**).

**Figure 4.**
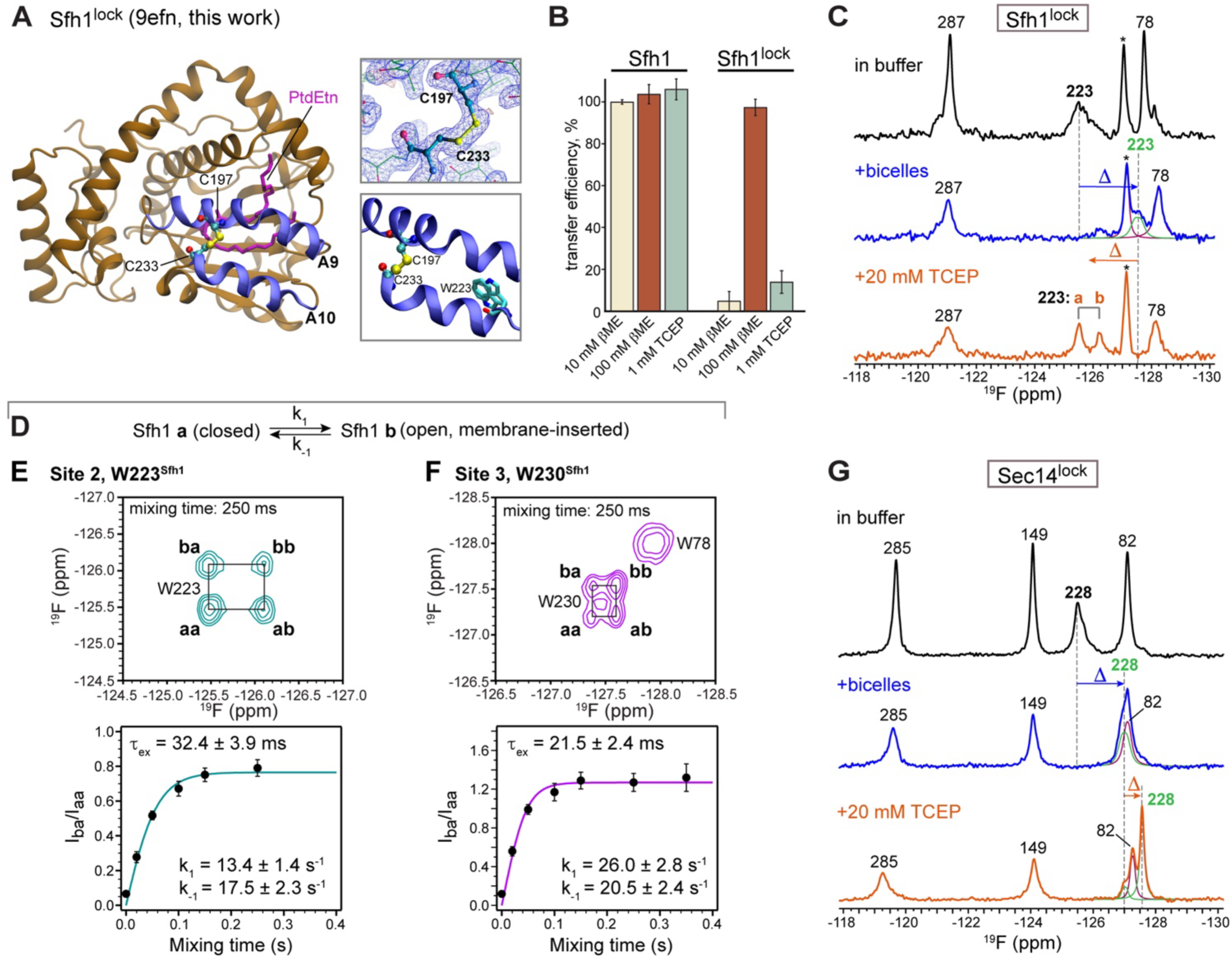
Sfh1/Sec14 gate dynamics in the membrane environment. **(A)** Crystal structure of the Sfh1^lock^ (Sfh1^F223W,K197C,F233C^) variant (PDB 9EFN, this work) and an expansion of the gate region showing the engineered disulfide bond and its position relative to the F^223^W substitution. The 2(F_o_-F_c_) electron density map is contoured at 1.5σ. **(B)** PtdIns transfer assays conducted for wild-type Sfh1 and Sfh1^lock^ in the absence and presence of reducing agents. Data represent the averages and standard deviations from 3 independent experiments involving the indicated conditions and compared to wild-type transfer values set at 100%. **(C)** ¹⁹F NMR spectra of Sfh1^lock^ in buffer (black) upon addition of bicelles (blue) and subsequent addition of 20 mM TCEP (orange). Reduction of disulfide bond in Sfh1^lock^ by TCEP results in recovery of two W_223_ conformers ‘a’ and ‘b’ observed in the wild-type protein. Peak labeled with (*) is a putative adduct of Cys thiol group and 5-fluoro-Trp. **(D)** Interconversion of gate conformers ‘a’ and ‘b’ where k_1_ and k_-1_ are the forward and reverse kinetic rate constants, respectively. ‘b’ is assigned to the open gate conformer based on the wild-type PRE data of Figure 3. **(E,F)** Two-dimensional [¹⁹F,¹⁹F] EXSY spectra at mixing time 250 ms show the presence of cross-peaks between the ‘a’ and ‘b’ conformers of Site 2 (**E**, W_223_^Sfh1^/green) and Site 3 (**F**, W_230_^Sfh1^/purple). The mixing time dependence of the intensity ratios of cross-peak (‘b’→‘a’) to auto-peak (‘a’→‘a’) was fit as described in Methods to obtain k_1_, k_-1_ and calculate τ_ex_=1/(k_1_+k_-1_). **(G)** ¹⁹F NMR spectra of Sec14^lock^ variant (Sec14^F228W,R195C,F231C^) in buffer (black) upon addition of bicelles (blue) and subsequent addition of 20 mM TCEP (orange). Reduction of disulfide bond in Sec14^lock^ by TCEP results in the recovery of the sharp W_228_ peak observed in the wild-type protein. Circa 20% of the molecules maintain the disulfide bond under these conditions as indicated by a minor peak at −127 ppm. Spectra containing closely spaced peaks in (**C**) and (**G**) were deconvoluted using Line Fitting tool in MestReNova; deconvoluted peaks of W_223_^Sfh1^ and W_228_^Sec14^ are in green.

The ^19^F NMR spectrum of 5-^19^F-W-Sfh1^lock^ in buffer shows three expected signals: two native Trp residues (W_287_ and W_78_) and the Site 2 probe W_223_ (**Fig. 4C**, black spectrum). There is an additional ^19^F peak. Mass-spectrometry data suggest this peak corresponds to an Sfh1 sub-population with an intramolecular W_287_-C_164_ cross-link.(Datta et al., 2001) Site 2 aromatic sidechain retains its mobility in the Sfh1^lock^ mutant as evidenced by the exchange-broadening of the ^19^F Sfh1^W223[lock]^ peak. The presence of two W_223_ sidechain rotamers in the Sfh1^lock^ crystal structure (**Fig. 4A**) suggests that the NMR-detected motions reflect aromatic ring flips.

Upon bicelle addition to the Sfh1^lock^ sample, the native ^19^F-W_287_ and ^19^F-W_78_ peaks broaden and the ^19^F-W_223_ signal shifts upfield by 2 ppm (**Fig. 4C**, blue spectrum). Subsequent addition of 20 mM TCEP results in the downfield shift of the ^19^F-W^223^ peak and the appearance of two ^19^F-W^223^ conformers ‘a’ and ‘b’ at the same chemical shifts observed in the sample of bicelle-associated ^19^F-W-substituted Sfh1^F223W^ (**Fig. 4C**, orange spectrum; **Fig. 3G**). In aggregate, these data indicate that: (i) the closed Sfh1 form is competent for membrane binding, and (ii) ^19^F-W_223_ conformers ‘a’ and ‘b’ are associated with A10 motions that open the gate. Since only peak ‘b’ has non-zero PRE (**Fig. 3G**), we assign ‘b’ to the open conformer where A10 inserts into the membrane, and ‘a’ to the closed conformer where A10 peripherally interacts with the membrane.

To determine the timescale of the ‘a’ to ‘b’ interconversion, we conducted two-dimensional [^19^F,^19^F] exchange spectroscopy (2D EXSY)(Palmer III et al., 2001) experiments on the Site 2 and Site 3 Sfh1 mutants. The timescale of the reaction τ_ex_ is defined as an inverse of the sum of the forward and reverse rate constants, k_1_ + k_-1_ (**Fig. 4D**). The interconversion between the open and closed Sfh1 conformers is evident from the appearance of the ‘ab’ and ‘ba’ cross-peaks for both ^19^F-W_223_ ^Sfh1^ and ^19^F-W_230_^Sfh1^ resonances (**Fig. 4E,F**). The rate constants k_1_ and k_-1_ were obtained from fitting the time dependence of peak intensity ratios (see Methods). The τ_ex_ values for ^19^F-W_223_^Sfh1^ and ^19^F-W_230_ ^Sfh1^ are comparable (32.4 ± 3.9 ms and 21.5 ± 2.4 ms, respectively; **Fig. 4E,F**). The slow dynamics of A10 opening and closing correlate well with Sfh1 being a slow lipid exchanger.

The same gate-locking strategy was applied to Sec14 to determine if the increase of Site 3 mobility upon membrane association reports gate motions. The cross-linked state of Sec14^F228W,R195C,F231C^ (Sec14^lock^) shows the expected three ^19^F signals with the ^19^F-W_228_^Sec14[lock]^ peak appearing at the same chemical shift of −125.5 ppm (**Fig. 4G**, black spectrum) as the ^19^F-W228^Sec14^ peak in the Site 3 Sec14 in buffer (**Fig. 3H**). Like Sfh1, the closed Sec14 conformer is competent to bind membranes. This property is evidenced by broadening of the native ^19^F-W149^Sec14[lock]^ and ^19^F-W285^Sec14[lock]^ peaks upon addition of bicelles. Bicelle addition also causes a 1.5 ppm upfield shift of the ^19^F-W_228_ peak based on deconvolution analysis (**Fig. 4G**, blue spectrum). However, it is only upon ‘unlocking’ the gate by addition of reductant that the ^19^F-W_228_^Sec14[lock]^ peak sharpens significantly due to the increase of the sidechain mobility (**Fig. 4G**, orange spectrum). The linewidth and chemical shift (−127.6 ppm) are comparable to those of the Site 3 reporter in the ^19^F-W-substituted Sec14^F228W^ spectrum (**Fig. 3H**). These results indicate that significant flexibility of the W_228_^Sec14^ sidechain is a signature of gate opening in membrane-bound Sec14. Based on the molecular mass of the protein-bicelle complex, we estimate the local correlation time of W_228_^Sec14^ sidechain motions is < 100 ns.

### Molecular dynamics simulations of the lipid exchange reaction

Experimental data produced the following conclusions: (i) closed conformers of Sec14/Sfh1 complexed to their native lipid ligand are dominant in solution; (ii) closed Sec14/Sfh1 conformers are competent for membrane binding; (iii) helix A10 is an essential membrane-inserting protein element; and (iv) membrane association triggers A10 dynamics that opens the gate and enables the lipid exchange reaction to occur. To arrive at the atomistic description of these events and gain insight into the mechanism of heterotypic lipid exchange, we carried out unbiased MD simulations of Sfh1 and Sec14 in the presence of membranes. PtdCho complexes of Sec14 (9efp) and Sfh1 (3b7q) were used as starting structures in systems with 1-palmitoyl-2-oleoyl-sn-glycero-3-phosphocholine/phosphoinositol/phosphoethanolamine bilayers (POPC/POPI/POPE=70:20:10, 128 total lipid molecules per leaflet). This composition mimics the proportions of these lipids in bulk yeast membranes. POPE was included as it enhances Sec14 lipid exchange activity in vitro.(Sugiura et al., 2021) Each protein was positioned with its center-of-mass 90 Å from the bilayer center (**Fig. 5A**). To generate four independent replicas, four different orientations relative to the membrane surface were applied to the protein molecule (**Table S3A**). The total duration of production runs was 8 μs each for Sec14 and Sfh1. The protein fold remained stable throughout the simulations, and membrane density profiles showed no perturbations of lipid distributions (**Fig. S5A,B**).

**Figure 5.**
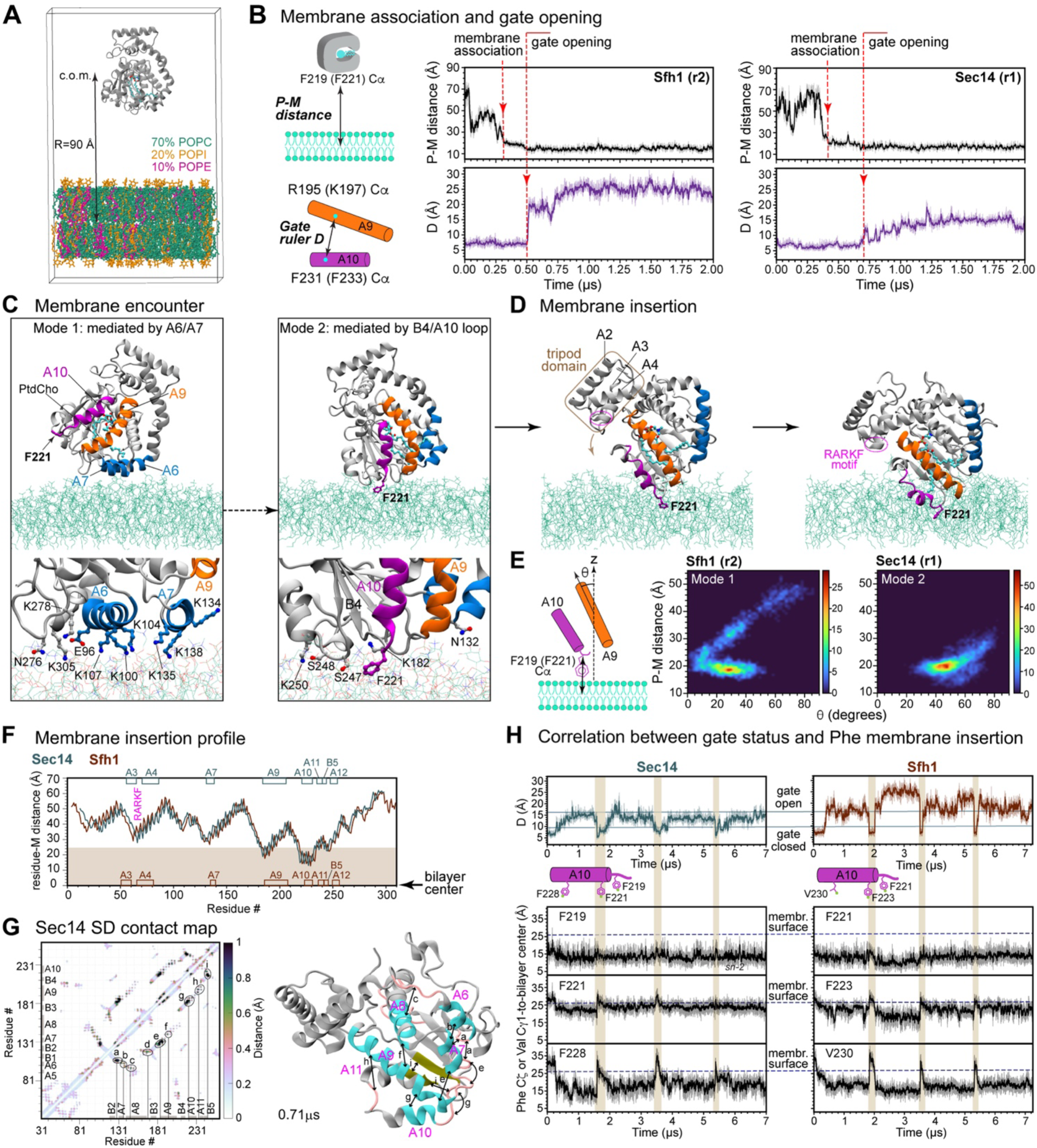
Sec14/Sfh1 membrane encounter/insertion and gate opening in MD simulations. **(A)** Composition and geometry of the simulated protein-membrane system. c.o.m. stands for center-of-mass. **(B)** Definitions of protein-to-membrane (P-M) and gate ruler (D) distances. Distances are plotted as a function of trajectory time to illustrate the relative timing of membrane engagement and gate opening. Both events are marked with red arrows. Replicas 1 (r1) and 2 (r2) are shown as representative data for Sec14 and Sfh1, respectively. **(C)** Two modes of membrane encounter, Mode 1 and Mode 2 detected in MD simulations and illustrated using the Sfh1 r2 data. Expansions of the protein-membrane interface highlight the residues involved in interactions with membrane lipids. **(D)** Illustration of the membrane insertion process A9 led by the F_221_ (Sfh1)/F_219_ (Sec14)-bearing loop and concludes with full insertion of A10 and the A9 N-terminal region. The RARKF motif (magenta) that is essential for PtdIns transfer activity is brought into proximity of the headgroup region. **(E)** Visualization of membrane encounter conformations using the tilt angle of helix A9 relative to the membrane normal and the P-M distance. **(F)** Residue-specific Sec14/Sfh1 membrane insertion profiles reported by the distance between the Cα atom and the bilayer center. The cytoplasmic leaflet of the bilayer is in beige. **(G)** Map of the r.m.s.d. values of inter-residue contacts in the membrane-bound Sec14. Secondary structure elements (α-helices, cyan; β-strands, olive; loops, salmon) that experience relative pairwise fluctuations due to the motions of either one or both elements are marked with black arrows. **(H)** Correlation between gate status and membrane insertion of NMR-characterized protein sites: loop residue Sec14 F_219_/Sfh1 F_221_ and A10 residues (Sec14 F_221_,F_223_,F_228_/Sfh1 F_221_,F_223_,V_230_) in concatenated trajectories of membrane-bound Sec14/Sfh1. Gate status is defined as open (D ≥ 16 Å) and closed (D < 10 Å). Intervals where the gate is closed are highlighted in beige.

### PITP membrane encounter and insertion

In all production runs, Sec14/Sfh1 spontaneously associated with the bilayer after several hundreds of nanoseconds and remained membrane-bound throughout the simulations. This is demonstrated using the distance between the Cα carbon of F_219_^Sec14^/F_221_^Sfh1^ and the center of the membrane bilayer (**Fig. 5B**). In agreement with our experimental data, A9-A10 gate opening occurs only after Sec14/Sfh1 are stably associated with membranes.

We discerned two productive membrane encounter modes (**Fig. 5C**). Mode 1 is primarily mediated by lysine ridges on helices A6 and A7 that engage in electrostatic interactions with lipid headgroups. Mode 2 involves peripheral interactions of F_219_^Sec14^/F_221_^Sfh1^ and the surrounding polar residues with the headgroup region. Both modes were present in the Sfh1 and Sec14 simulations and, in all cases, protein bound in mode 1 reoriented to assume mode 2. Subsequently, Sec14/Sfh1 insert deeper into the membrane and undergo a rotational motion. This process leads to the A10 and N-terminal end of A9 penetrating into acyl chain region of the membrane, and to the positioning of the RARKF motif near the membrane surface (**Fig. 5D**). The RARKF motif resides on the tripod domain of Sec14-like PITPs and is essential for PtdIns headgroup binding.(Phillips et al., 1999; Schaaf *et al*., 2008) **Video S1** summarizes the action from initial membrane encounter to opening of the A9/A10 gate.

A holistic view of the combined membrane encounter and insertion process is shown in representative heat maps that correlate the distance between F_219_^Sec14^/F_221_^Sfh1^ Cα and the bilayer center to Sec14/Sfh1 orientation relative to the membrane surface (**Fig. 5E**). The latter parameter is reported as the tilt angle between A9 and the membrane normal. Irrespective of initial encounter mode, Sec14/Sfh1 converge on very similar membrane-bound states characterized by a distance of ∼ 19 Å and tilt angles of 30-60 degrees. To quantitatively compare residue-specific membrane insertion profiles between Sec14 and Sfh1, the trajectories of the bilayer-associated Sec14/Sfh1 were concatenated to yield a >7 μs combined trajectory for each. A plot of mean distances between the residue c.o.m. and the bilayer center identified the Sec14/Sfh1 structural elements that engage with membranes (**Fig. 5F**). These elements are: (i) the N-terminal segment of A9, and helices A10/A11 that reach into the acyl region; (ii) the B5-A12 segment that peripherally interacts with the headgroup region; and (iii) the A3/A4 turn and helix A7 that are positioned above the surface but can engage protruding lipid headgroups (**Fig. S5C**). These data support our conclusions from experiments that highlight A10 as the major membrane insertion element and identify other protein regions involved in stable membrane recruitment.

### Conformational plasticity of membrane-bound Sec14-like PITPs

Backbone r.m.s.f. profiles identify the A9/A10 gate as the most dynamic element of the membrane-bound Sec14 PITPs with A10 consistently showing the largest r.m.s.f. values in all simulations (**Fig S5D**). Other protein elements with elevated dynamics are helix A7 and the A3/A4 turn that harbors the RARKF PtdIns recognition motif. To evaluate the effect of these motions on 3D protein structure, we constructed a map of the r.m.s.d. values of inter-residue contacts along the concatenated trajectories of membrane-bound Sec14 and Sfh1 (**Fig. 5G**, **S5G**). These analyses identified secondary structure elements that experience relative pairwise fluctuations due to the motions of either one or both elements. The dynamic patterns of Sec14 and Sfh1 are similar and involve helices A6/A7/A8/A9/A10, β strands B5/B4 and the adjacent loop regions (**Fig. 5G**, **S5G**). As all of these elements frame the lipid-binding pocket of Sec14-like PITPs, their relative motions directly report on the conformational plasticity of the bound lipid’s environment.

To determine if the MD simulations are consistent with experimental conclusions, we examined the relationship between the gate status of membrane-bound Sec14 and Sfh1 and insertion depth of the A10 residues that were probed by ^19^F NMR spectroscopy. In both proteins, F_219_^Sec14^/F_221_^Sfh1^ stably insert into the bilayer prior to gate opening and remain anchored when the gate opens (**Fig. 5H**). By contrast, membrane insertion of F_221_^Sec14^/F_223_^Sfh1^ and F_228_^Sec14^/V_230_^Sfh1^ is tightly correlated with the gate opening event (**Fig. 5H**). Notably, F_221_^Sec14^ does not insert deeply into the bilayer. This behavior accounts for the absence of a PRE effect at this site in ^19^F-NMR experiments (**Fig. 3F**).

### Sec14/Sfh1 demix PtdIns in the membrane

Lipid density plot analyses of the concatenated trajectories reveal that both membrane-associated Sec14 and Sfh1 collect (or demix) PtdIns molecules along the periphery of the protein-membrane interface (**Fig. 6A**). Demixing is mediated by protein residues whose side chains serve as H-bond donors or acceptors for the hydroxyls of the inositol ring. In addition, opportunistic salt bridges are formed between the PtdIns phosphoester group and Lys/Arg sidechains that reach into the headgroup region. The same set of secondary structure elements in Sec14/Sfh1 is involved in PtdIns recruitment (**Fig. 6A**). For Sec14, these regions belong to helices A9: _195_<u>R</u>EASYI<u>SQ</u>_202_, A10: _222_<u>S</u>TAF<u>R</u>LF<u>K</u>_229_, A12: _245_<u>SS</u>Y<u>QKE</u>LL<u>K</u>_253_, and the A3/A4 turn that harbors the _63_RA<u>RK</u>F_67_ PtdIns recognition motif. Sec14/Sfh1 also induce demixing of PtdEtn molecules, where the lipid amine group forms salt bridges with negatively charged Sec14/Sfh1residues (including D_233_/D_235_). Formation of this salt bridge might facilitate gate opening since D_233_/D_235_ stabilize Sec14/Sfh1 closed gate conformations in membrane-free systems via interactions with Q_202_/Q_204_ (**Fig. S6A**). The PtdIns and PtdEtn clustering regions overlap near A9 and the tripod motif but not anywhere else. Overall, PtdIns clustering around Sec14/Sfh1 not only increases its local concentration but also enables PtdIns recruitment via coordination of the inositol head group by the RARKF motif of the tripod domain.

**Figure 6.**
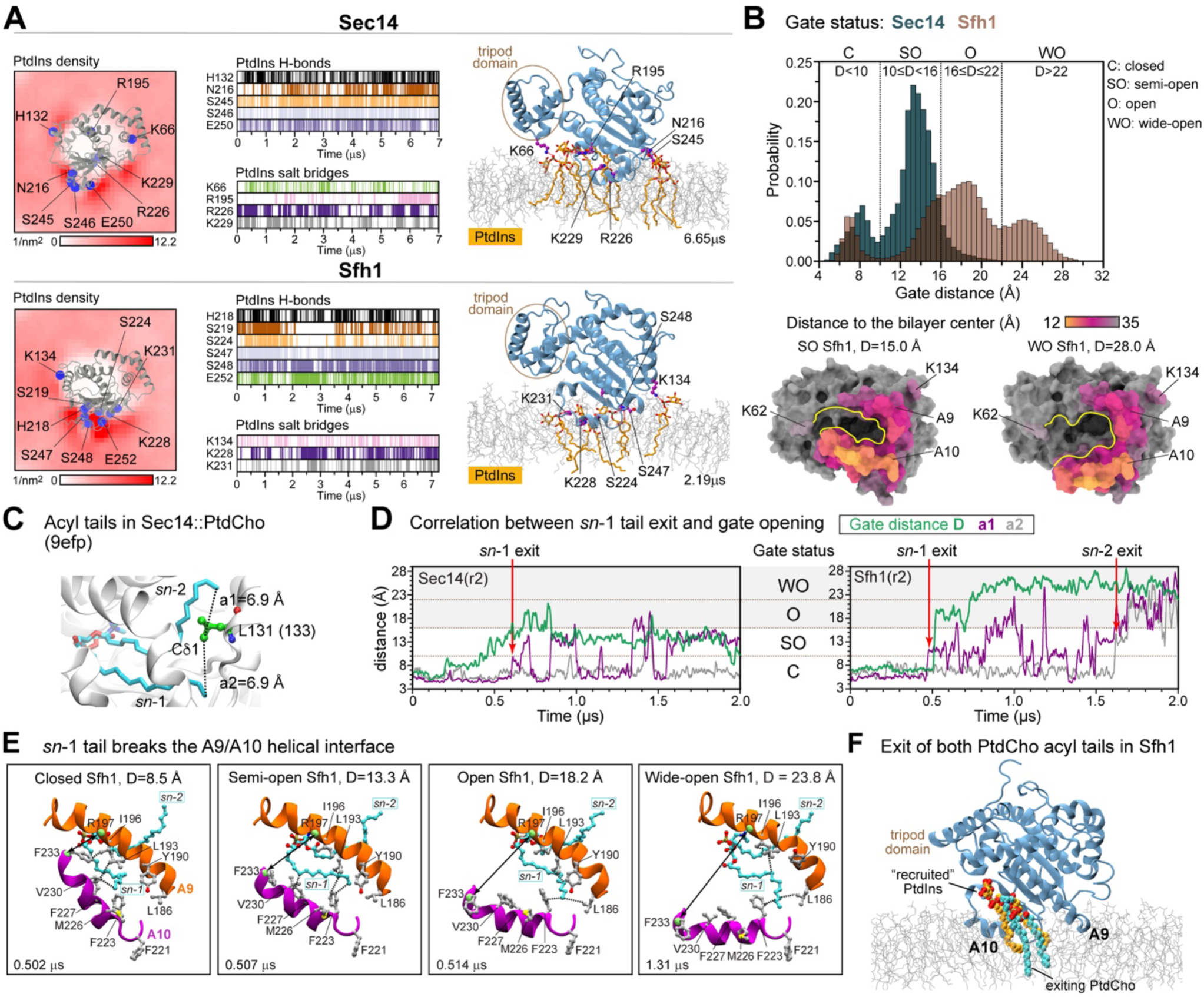
Phospholipid interactions of membrane-associated Sec14/Sfh1 and conformational dynamics of the gate. **(A)** Demixing of PtdIns by Sec14/Sfh1 illustrated by two-dimensional lipid density plots, event plots for the most populated protein-PtdIns H-bonds and salt bridges, and representative MD snapshots (Sec14/Sfh1-bound PtdCho is omitted for clarity). **(B)** Histogram of gate distance distributions in membrane-associated Sec14 (blue) and Sfh1 (brown). The gate states are defined as closed (D < 10 Å), semi-open (10 ≤ D < 16 Å), open (16 ≤ D ≤ 22 Å) and wide open (D < 22 Å). Residue-to-bilayer center distances are color-coded and mapped onto the surface representations of semi-open (SO) and (WO) Sfh1 conformers. Gate mouth is outlined in yellow; bound PtdCho is omitted for clarity. **(C)** PtdCho pose in the lipid-binding Sec14 pocket, parameterized by the distances a1 and a2 between the Cδ1 of Sec14 L_133_ and methyl carbons of *sn*-1 and *sn*-2, respectively. **(D)** Gate ruler D, a1 and a2 plotted as a function of time for Sec14 r2 (left) and Sfh1 r2 (right). Points of acyl chain *sn*-1 and *sn*-2 exit from the lipid binding pocket are indicated by arrows. **(E)** Snapshots from the Sfh1 r2 production run depicting the gate region defined by helices A9 (orange)/A10 (violet) and the pose of PtdCho (cyan). The respective gate ruler distances D are shown with black arrows. Key Sfh1 residues are in gray. **(F)** The snapshot from the Sfh1 r2 production run used as a starting structure for replicas 5, 6, 7, and 8 (see Methods).

### Sec14/Sfh1 gate opening correlates with the exit of the sn-1 acyl chain of bound PtdCho

Analysis of gate distance (D) distributions in membrane-bound Sec14 and Sfh1 revealed a broad range of values we group into four categories according to gate status: closed (D < 10 Å), semi-open (10 ≤ D < 16 Å), open (16 ≤ D ≤ 22 Å), and wide-open (D > 22 Å) (**Fig. 6B**). Membrane-bound Sec14 showed a strong preference for semi-open conformers on the MD simulation timescales. Sfh1 adopted a broader range of D values with significant sampling of wide-open conformations with D values reaching *ca.* 30 Å (**Fig. 6B**). While gate opening drastically increases membrane exposure of the lipid-binding pocket it does not significantly influence depth of membrane insertion or its orientation (**Fig. 6B**, **S6B,C**).

To describe the behavior of bound PtdCho during the gate opening, the PtdCho pose in the lipid-binding pocket was parameterized as the distance between the terminal methyl carbon of the *sn*-1 and *sn*-2 acyl tails and the Cδ1 carbon of L_131_^Sec14^/L_133_^Sfh1^ (**Fig. 6C**). We find that exit of the *sn*-1 acyl chain from the binding pocket (manifested as increase in the corresponding distance) correlates with the gate adopting an open conformation with D ≥ 16 Å (**Figs. 6D, S6D**). The mechanism by which bound PtdCho facilitates gate opening is through intercalation of its *sn*-1 acyl chain between helices A9 and A10 (**Fig. 6E**). The *sn*-1 acyl tail progressively invades the A9/A10 interface by forming van der Waals contacts with hydrophobic A9 residues, while the ‘liberated’ A10 helix swings open into the bilayer acyl chain environment. These findings suggest the dynamics of bound PtdCho plays a key role in initiating the transition of Sec14/Sfh1 from closed to open conformers.

Of the eight replica simulations, only two Sfh1 production runs produced significant movement of bound PtdCho as defined by both *sn*-1 and *sn*-2 acyl tails exiting the binding pocket. Using this configuration as initial structure (**Fig. 6F**), four independent 2.5 μs replica simulations were conducted. In all four replicas, PtdIns replaced PtdCho in the lipid binding pocket (**Fig. S6E**). Analysis of these trajectories revealed the key features of the lipid exchange process.

### The nuts and bolts of heterotypic PtdCho/PtdIns exchange process

Lipid exchange is initiated by gate opening – an action facilitated by egress of the PtdCho *sn*-1 acyl chain from the lipid binding pocket. The exited *sn*-1 acyl moiety then moves freely within the open gate region where it forms transient van der Waals contacts with hydrophobic residues of helices A9/A10 and dips its distal region into the bilayer. The protein context in which PtdCho resides at this stage is as follows. The choline headgroup is enveloped in an aromatic cage (Y_113_, Y_124_, Y_153_, F_156_) while the non-ester oxygens of the phosphoester moiety are situated in a polar environment and are in H-bond contact with hydroxyl-carrying sidechains (Y_153_, Ser_175_, T_177_). The *sn*-1 and *sn*-2 acyl chain carbonyl oxygens engage in H-bond interactions with the sidechains of S_203_Q_204_ (**Fig. 7A**, panel 1; **Fig. S7Ai**).

**Figure 7.**
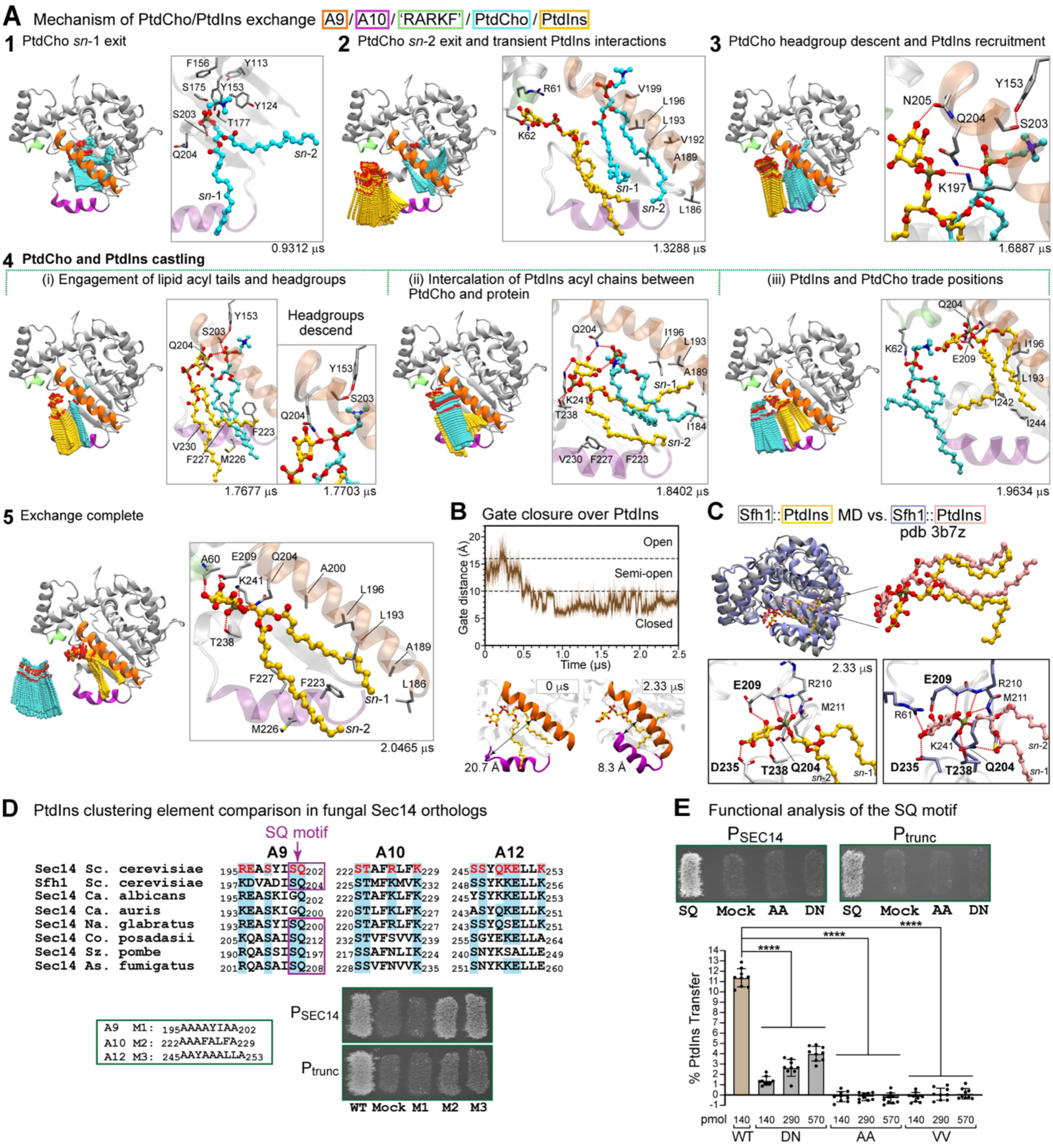
PtdCho/PtdIns exchange process. **(A)** Five general stages of the PtdCho/PtdIns exchange. For each stage, left panels show a composite view of PtdCho (cyan) and PtdIns (gold) positions as the exchange reaction progresses through the indicated stages. Right panels show representative snapshots of the respective lipid poses. Stage 4 (“castling”) is subdivided in three steps i through iii. **(B)** Gate distance plotted against time depicts closing of the Sfh1 gate over newly bound PtdIns. A close-up view of gate closing is shown as comparison of the A9 (orange)/A10 (purple) poses at the beginning and end of the simulation. **(C)** Top panel: backbone superposition of the simulated closed Sfh1::PtdIns conformer at 2.33 μs (gray/gold) and the Sfh1::PtdIns crystal structure (purple/pink). The respective poses of bound PtdIns are at right. Bottom panel: PtdIns headgroup coordination in the simulated closed Sfh1::PtdIns conformer (left) and Sfh1::PtdIns crystal structure (right). **(D)** Top panel: alignment of the PtdIns-clustering element sequences identified for Sec14/Sfh1 with those of Sec14 PITPs from the indicated *Candida*, *Nakaseomyces*, *Coccidioides*, *Schizosaccharomyces*, and *Aspergillus* species. Sequence identities are highlighted in blue, and *S. cerevisiae* Sec14 residues substituted with Ala in the Ala-scan experiment for each element are in red. Bottom panel: fatal attraction data for the Ala-scan mutants M1, M2, and M3 are shown under conditions of expression from the native *SEC14* and truncated promoters (P^SEC14^, P_trunc_, respectively) as indicated. **(E)** Functional role of SQ motif. Top panel: fatal attraction assay results for the indicated Sec14 variants expressed under the control of the native and truncated promoters. Bottom panel: PtdIns-transfer activities of the indicated purified Sec14 variants relative to that of the wild-type protein.

Subsequent exit of the PtdCho *sn*-2 acyl chain from the lipid binding pocket is promoted by its Van der Waals interactions with hydrophobic sidechains of A9. While individual interactions are transient, the involvements of multiple *sn*-2 methylene groups align the PtdCho acyl chain along the hydrophobic face of A9 (**Fig. 7A**, panel 2). In the meantime, the basic residues of the tripod domain RARKF motif(Phillips *et al*., 1999; Schaaf *et al*., 2008) bring PtdIns molecules in close proximity to the gate mouth by forming H-bonds with the inositol ring hydroxyls and salt bridges with the non-ester phosphate oxygens.

The next stage of PtdCho/PtdIns exchange is marked by the persistent recruitment of the incoming PtdIns to the open gate mouth and descent of the PtdCho headgroup from the pocket interior towards the membrane bilayer. PtdCho headgroup descent comes with the choline group leaving the aromatic cage, disruption of the Y_153_-phosphodiester H-bond, and replacement of that H-bond with one involving the sidechain of Q_204_ (**Fig. 7A**, panel 3; **Fig. S7Ai**). The PtdIns headgroup also shifts into closer proximity to the gate by engaging with hydrophilic residues at the C-terminal end of A9 (K_197_, N_205_; **Fig. 7A**, panel 3; **Fig. S7Aii**). These two actions bring the positions of the PtdCho/PtdIns headgroups to level and set the phospholipids up for a critical repositioning we define as “castling”.

Initiation of castling involves engagements of PtdCho acyl chains with those of PtdIns in Van der Waals interactions and H-bond interactions between the PtdIns headgroup hydroxyls and the PtdCho phosphate oxygens (**Fig 7A**, panel 4i; **Fig. S7Aiii**). The former are facilitated by a hydrophobic ‘rail’ provided by helix A10 (F_223_, M_226_, F_227_, V_230_). The latter takes advantage of A9 residues S_203_ and Q_204_ whose sidechains form H-bonds with the PtdCho non-ester phosphate oxygen and the *sn*-2 carbonyl oxygen, respectively. Both headgroups subsequently drop toward the membrane surface in a concerted action where Q_204_ engages with PtdIns while PtdCho loses its H-bond interaction with S_203_ (**Fig. 7A**, panel 4i). Subsequently, the PtdIns acyl chains position themselves between the PtdCho chains and A10 with the net result that PtdCho is displaced from the hydrophobic A10 rail (**Fig. 7A**, panel 4ii). The final stage of castling completes the reciprocal trade of PtdCho and PtdIns positions relative to the lipid binding pocket. The exiting PtdCho is largely displaced from the gate mouth and remains only loosely associated to protein by transient interactions with Q_204_ and K_62_ of the RARKF motif (**Fig. 7A**, panel 4iii; **Fig. S7Ai**) and with hydrophobic residues of the C-terminal half of A10. PtdIns fully occupies the gate mouth at this point. The PtdIns acyl chains are involved in van der Waals contacts with hydrophobic residues of A9 and strand B5 that is an element of the lipid pocket ‘floor’, and the inositol headgroup is in H-bond contact with the sidechains of Q_204_ and E_209_ (**Fig. 7A**, panel 4iii; **Fig. S7Aii**).

After castling is complete, PtdIns is positioned between residues Q_204_/K_62_ (**Fig. S7B**) and PtdCho interactions with these residues and with A10 break -- resulting in the phospholipid diffusing away into the larger membrane surface (**Fig. S7C**). The bound PtdIns headgroup remains held at the open gate via engagement with Q_204_, E_209_, T_238_, K_241_ – i.e. residues that form the PtdIns headgroup binding network in the Sfh1::PtdIns crystal structure. RARKF motif residues K_62_ and A_60_ also contribute. The *sn-1* acyl chain is packed against the hydrophobic face of A9, and the *sn-2* chain is engaged with the A10 hydrophobic rail (F_223_, M_226_, F_227_) (**Fig. S7A**, panel 5). A movie summarizing the PtdCho/PtdIns exchange reaction is provided in **Video S2**.

### The post-exchange fate of PtdIns

Additional simulations of the membrane-bound Sfh1::PtdIns complex revealed that PtdIns remained bound to the protein at the gate mouth and occasionally penetrated deeper into the binding pocket interior. However, PtdIns did not fully incorporate into the lipid binding pocket, and, other than one very transient instance, the gate did not progress past semi-open states to closed states in any of these simulations. To test the fitness of the Sfh1::PtdIns complex formed in the membrane context, that structure was used as initial configuration to conduct 2.5 μs MD simulations in an aqueous membrane-free system. The gate transitioned to a closed (D<10 Å) conformation at *ca.* 500 ns (**Fig. 7B**). Analysis of the gate distance distributions shows a clear conformational preference with D values of ca. 7 and 9 Å (**Fig. S7D**). The PtdIns poses in closed MD-derived conformers approximate the one observed in the Sfh1::PtdIns crystal structure (**Fig. 7C**). The most significant difference is in the positions of the acyl chains that maintain considerable flexibility throughout the simulations (**Video S3**). The PtdIns headgroup adopts a pose similar to that described in the Sfh1::PtdIns crystal structure. It resides in a polar environment where the Q_204_, E_209_, D_235_, T_238_, and K_241_ sidechains interact with the inositol ring hydroxyls and oxygens of the phosphate moiety (**Fig. 7C**).

### Biological relevance of newly recognized Sec14 functional elements

An important feature of the MD simulations is a functional assignment for two classes of previously uncharacterized motifs in Sec14 and Sec14-like PITPs. The first is defined by the elements involved in PtdIns H-bond interactions and PtdIns demixing at the Sec14/Sfh1-membrane interface -- _195_<u>R</u>EASYI<u>SQ</u>_202_ (helix A9), _222_<u>S</u>TAF<u>R</u>LF<u>K</u>_229_ (helix A10), and _245_<u>SS</u>Y<u>QKE</u>LL<u>K</u>_253_ (helix A12). These three elements (termed M1, M2, M3, respectively) are conserved in Sec14 orthologs of highly diverse fungi that span vast evolutionary distances. This conservation suggests PtdIns demixing is an intrinsic property of Sec14-like PITPs (**Fig. 7D**, top panel). Alanine-scanning mutagenesis experimentally assessed the physiological importance of each individual element in two versions of a fatal attraction assay. The first is a native context where mutants are expressed from a full-length *SEC14* promoter. This native level of expression is at least 10-fold in excess over what is required for vegetative cell growth and can phenotypically obscure significant functional defects in mutant Sec14 proteins (Salama et al., 1990; Phillips et al., 1999; Nile et al., 2014). The second is a ‘sensitized’ context that reveals partial defects when mutants are expressed at a reduced level (i.e. from a truncated promoter) sufficient to meet the Sec14 functional threshold required for cell viability. Reconstitution of cells with a Sec14 altered for M1 as sole source of Sec14 activity fails both versions of the complementation test whereas reconstitution with M2 or M3 supports viability in both – albeit very poorly in the sensitized context (**Fig. 7D**, bottom panel). All three Sec14 variants are stable proteins in vivo (**Fig. S7E**, top panel). We interpret these results to indicate that the PtdIns clustering elements each contribute to Sec14 activity in vivo with M2 and M3 exhibiting a measure of functional redundancy.

The role of element M1 cannot be interpreted solely in terms of PtdIns demixing as it includes the SQ motif. That is, the second class of uncharacterized motifs in Sec14 and Sec14-like PITPs projected to be key for PtdCho exit from the lipid binding pocket, in PtdIns/PtdCho castling and in PtdIns entry into the lipid binding pocket. Again, the SQ motif is conserved in Sec14 PtdIns/PtdCho exchange proteins of highly divergent fungi that span vast evolutionary distances (**Fig. 7D**, top panel). Although the Sec14^SQ201,202AA^ mutant is stable in vivo (**Fig. S7E**, bottom panel), it scores as defective in both native and sensitized contexts of the ‘fatal attraction’ complementation test (**Fig. 7E**, top panel). The Sec14^SQ201,202DN^ mutant is also a stable protein in vivo (**Fig. S7E**, bottom panel), and it too fails both versions of the complementation test. These in vivo defects were recapitulated in vitro. Purified Sec14^SQ201,202AA^ and Sec14^SQ201,202DN^ are strongly defective for PtdIns-transfer activity (**Fig. 7E**; bottom panel) – even though circular dichroism spectra report both to be properly folded proteins (**Fig. S7F**). Sec14^SQ201,202VV^ is unstable when expressed in yeast but purifies from *E. coli* as a well-folded protein (**Fig. S7F**). Sec14^SQ201,202VV^ is also inactive for PtdIns-transfer in vitro (**Fig. 7G**). These results identify the SQ motif as a critical element required for the biological activities of Sec14 and of its orthologs.

## DISCUSSION

A principal basis for homeostatic cell control relies on the utility of peripheral membrane proteins as core regulators of membrane-based signal transduction systems. Phosphoinositide signaling is a major pillar of such homeostatic control. The PtdIns 4-OH kinase/PITP partnership required for PtdIns4P signaling represents a unique arrangement where two peripheral membrane proteins must cooperate to produce biologically relevant PtdIns4P pools. The dominant PITP partners in the *Eukaryota* are Sec14, Sec14-like PITPs and related yeast, plant, insect and mammalian proteins of the Sec14/CRAL-Trio fold – i.e. a highly conserved and highly expanded module for binding lipids and other small hydrophobic molecules.(Aravind et al., 1999; Huang et al., 2016a; Montag *et al*., 2023; Saito et al., 2007)

Heterotypic PtdCho/PtdIns exchange is the essential engine by which Sec14-like PITPs potentiate the activities of PtdIns 4-OH kinases in vivo.(Bankaitis *et al*., 2025; Schaaf *et al*., 2008) Lipid exchange operates without additional energy input and involves lipids that both reside in favorable thermodynamic environments. The very nature of this process raises fundamental questions: (i) How do Sec14-like PITPs interface with membranes during lipid exchange? (ii) How does protein conformational dynamics enable lipid exchange? (iii) Can (does?) lipid exchange occur in a concerted manner, or is there an obligatory apo-PITP intermediate? (iv) Do the invading and exiting lipids interact to ensure a coupled lipid exit/entry mechanism? Herein, we address key outstanding questions and provide a framework for a mechanistic understanding of the Sec14/CRAL-Trio protein superfamily.

### Conformational plasticity of Sec14-like PITPs during lipid exchange

The major conformational transition in Sec14-like PITPs is the separation of helices A9 and A10 that gate entry into the lipid binding pocket. Evidence to this effect is provided both by crystal structures (Sec14::PtdCho and Sec14::PtdCho::NPPM481) and MD simulations. This transition is not required for membrane association as ^19^F NMR experiments with gate-locked mutants and MD simulations report this step is executed by the closed conformers that are dominant in solution. The MD simulations further show that gate opening ensues after the gate helices of closed Sec14::PtdCho and Sfh1::PtdCho conformers penetrate into the membrane bilayer (A_10_ deeply and A_9_ shallowly). That conclusion is buttressed by experimental evidence that includes: (i) ^19^F-NMR paramagnetic relaxation data demonstrating that the Sec14/Sfh1 A_10_ helices insert into the bilayer, and (ii) mutagenesis data consistent with A10 insertion being critical for Sec14 activity in vitro and in vivo. Neither Sec14 nor Sfh1 position the open gate mouth flat against the membrane surface -- irrespective of gate status (semi-open, open, or wide-open). Rather, these PITPs assume tilted poses that position the apex of the open gate mouth and the RARKF element of the tripod domain above the membrane surface.

Collectively, our experimental and computational data demonstrate that full gate opening – an essential step in lipid exchange– requires an action that breaks the A9/A10 interface. In the membrane environment, this action is facilitated by intercalation of the *sn*-1 acyl chain of exiting PtdCho between gate helices A9 and A10. An independent example of this concept is provided by the Sec14::NPPM481::PtdCho ternary complex formed in a membrane-free context. In this case, the small molecule inhibitor displaces resident PtdCho from its canonical pose in the lipid binding pocket with the result that the gate opens to accommodate the displaced PtdCho whose pose is now primarily stabilized by its hydrophobic interactions with A10. In addition to gate motions, the conformational plasticity of Sec14/Sfh1 is manifested in relative fluctuations of other secondary structure elements (helices A6/A7/A8, β strands B5/B4, and the adjacent loop regions) that frame the lipid binding pocket.

### Lipid demixing

Our MD data report Sec14/Sfh1 recruit PtdIns from the bulk lipid phase to specific points along the protein periphery. We suggest such a local PtdIns enrichment activity contributes to the Sec14/Sfh1 preference for PtdIns. This idea is reinforced by conservation of the three Sec14 motifs involved in this PtdIns demixing by its orthologs and the contributions of these apparently functionally redundant motifs to Sec14 activity in vivo. These data also provide a rationale for why PITPs exhibit a significant preference for PtdIns than for PtdCho.(Somerharju et al., 1983; Szolderits et al., 1989) Membrane-bound Sec14/Sfh1 also induce PtdEtn demixing from the bulk lipid phase. A functional relevance of this property is supported by the enhancement of Sec14 lipid exchange activity by PtdEtn in vitro.(Sugiura *et al*., 2021) The pattern of PtdEtn headgroup-protein H-bonds and salt bridges suggests PtdEtn might stabilize the open gate via interactions with hydrophilic residues of A9 and A10. Demixing of this non-bilayer-forming lipid might also disturb the local membrane environment at the PITP/bilayer boundary with the effect of lowering the energy barrier for lipid exchange.

### PtdCho exit

PtdCho exit from the lipid-binding pocket requires transition of the gate to a wide-open configuration. In that regard, Sec14 primarily adopts semi-open gate configurations in the MD simulations whereas Sfh1 shows a much higher preference for wide-open conformers. This behavior accounts for our inability to capture PtdCho/PtdIns exchange by Sec14. The gate helices A9 and A10 provide a scaffold that organizes PtdCho exit. Helix A9 accommodates the PtdCho headgroup as it navigates the hydrophobic lipid binding pocket from its initial pose to the open gate. The requisite interactions are provided by Y_153_ and the S_203_Q_204_ pair -- an essentially invariant feature of Sec14 PtdIns/PtdCho PITPs that resides at the C-terminus of helix A9. The hydrophobic face of A9, made accessible upon gate opening, successively engages *sn*-1 and *sn*-2 PtdCho acyl chains as these exit the pocket. In addition to being a dominant membrane insertion element, helix A10 also provides a hydrophobic rail along which the acyl chains of the exiting PtdCho and incoming PtdIns slide.

### Concerted nature of lipid exchange

We describe a heterotypic exchange reaction where the invading PtdIns first engages the RARKF motif of the tripod domain and is subsequently brought into the open gate mouth where it is positioned in immediate proximity of the exiting PtdCho. The most surprising outcome of our MD simulations is just how extensively the invading PtdIns and the exiting PtdCho interact in the concerted exchange reaction. These PtdIns-PtdCho interactions foster reciprocal trade of the phospholipid positions relative to the gate mouth. We posit this castling action defines the committed step in the exchange reaction. PtdIns recruitment to the open gate does not ensure exchange of PtdCho for PtdIns, however. PtdIns molecules are captured by RARKF in the open Sfh1 and semi-open conformations of Sec14 in multiple simulations. Yet, these PtdIns molecules fail to be exchanged for PtdCho and ultimately diffuse away into the bilayer. PtdIns must switch its engagement from RARKF to A9 to be positioned for interaction with the exiting PtdCho at the open Sec14/Sfh1 gate. This switch is required for PtdCho/PtdIns castling to occur with full exchange to follow.

### Post-exchange behavior of the system

We observed neither the closure of the gate over PtdIns that replaced the resident PtdCho nor the release of the PtdIns-loaded protein from the membrane on the timescale of our MD simulations. However, the Sfh1::PtdIns complex adopted a closed conformation (with gate distances fluctuating between 6 and 10 Å) when membrane was removed from the system. We attribute these gate fluctuations to the PtdIns acyl chains not yet having adopted their final poses using a high resolution Sfh1::PtdIns crystal structure as a reference for the final PtdIns-loaded state.

Our NMR experiments show bicelle-bound Sfh1 interconverts between open and closed states on the timescale of *ca.* 20-30 ms. This observation raises the possibility that Sec14/Sfh1 undergoes multiple rounds of lipid exchange on membranes prior to dissociating. In the biological context, such a scenario would require that the membrane residence time exceeds the timescale of the lipid-exchange reaction whose rate-limiting step could be the conformational dynamics of the gate element. This is not a concept considered in contemporary discussions of PITP function. Current views imply PITPs execute a single round of lipid exchange per membrane binding event.

### Implications for how Sec14-like PITPs potentiate PtdIns4P signaling in cells

The available data identify Sec14 as a PtdCho sensor that coordinates PtdCho metabolism with PtdIns4P signaling in late Golgi/early endosomal compartments.(Bankaitis *et al*., 2025) Sec14 enhances PtdIns 4-OH kinase activity in yeast -- even though PtdIns is a major (and in some strains the major) membrane phospholipid. Thus, PtdIns 4-OH kinases are intrinsically insufficient enzymes that must partner with a PITP to support productive PtdIns4P signaling in vivo. This is the case in yeast, plant and mammalian systems,(Huang et al., 2016b; Montag *et al*., 2023; Schaaf *et al*., 2008; Xie and Bankaitis, 2022) but the underlying mechanism is not understood. The MD simulations provide insights into how the Sec14 heterotypic lipid exchange reaction might make PtdIns a more accessible substrate for the enzyme.

The concerted PtdCho exit/PtdIns entry pathway we describe includes events where the exiting PtdCho interferes with PtdIns invasion of the lipid binding pocket. This ‘traffic jam’ transiently lifts the RARKF-associated PtdIns headgroup above the membrane surface – a pose that might enhance PtdIns accessibility to the PtdIns 4-OH kinase. This enhancement could come in one of two ways -- a direct channeling of that PtdIns molecule to the kinase, or a more indirect mechanism where Sec14 clusters PtdIns and provides a local substrate-rich environment for the enzyme. Regarding the latter idea, all three PtdIns clustering elements are preserved in mutants capable of only homotypic PtdIns/PtdIns exchange – yet these mutants are incapable of enhancing PtdIns 4-OH kinase activity in vivo. Thus, the most parsimonious interpretation of the data is that PtdIns 4-OH kinase enhancement requires PtdIns engagement with the gate mouth in the context of a PtdCho-bound open Sec14 conformer. This arrangement suggests PtdIns recruitment to the gate mouth has three potential outcomes: (i) completion of lipid exchange, (ii) biologically productive phosphorylation by PtdIns 4-OH kinase in a substrate channeling mode, or (iii) disengagement without exchange or phosphorylation. Only outcome (ii) consumes ATP.

### Limitations of the study

Key questions remain. First, we have yet to model Sec14/Sfh1 dissociation from the membrane. How Sec14-like PITPs are incentivized to execute this final step remains unresolved. Capturing this event will require longer simulation timescales and accelerated sampling MD approaches. Second, the mechanisms of homotypic PtdCho/PtdCho and PtdIns/PtdIns exhanges are likely to have unique features. This is of particular interest for homotypic PtdIns/PtdIns exchanges as these fail to enhance PtdIns 4-OH kinase activities in vivo. Finally, although Sfh1 is a faithful model for Sec14-mediated lipid exchange, why were we able to simulate lipid exchange for the slow exchange Sfh1 but not for the fast exchange Sec14? Perhaps Sfh1 has reduced affinity for PtdCho – a property that would favor its exit from its binding pocket in MD simulations. Sfh1* fast exchange mutants must hold the clues to this conundrum, but those clues are not yet understood.

## EXPERIMENTAL PROCEDURES

### Materials

All common chemicals such as buffer reagents and salts were purchased from Thermo Fisher Scientific or Millipore Sigma. 5-Fluoro-L-tryptophan was acquired from Millipore Sigma. Isopropyl-β-D-thiogalactopyranoside (IPTG) and Tris(2-carboxyethyl)phosphine hydrochloride (TCEP) were obtained from Research Products International. Chromatography columns and supplies were from Cytiva, and all lipids were from Avanti Research.

### Yeast strains, plasmids and media

Yeast strains included CTY303 (*MATa ura3-52 lys2-801 Δhis3-200 cki1-1 sec14Δ1::HIS3*) and CTY335 (MATα *ura3-52 Δhis3-200 sec14-129::HIS3 ade2-101 Δtrp1 kes1-124*).(Bankaitis *et al*., 1989; Cleves *et al*., 1991; Phillips *et al*., 1999) Yeast peptone dextrose (YPD) and minimal standard defined (SD) media have been described.(Sherman, 1983) Solid YPD and minimal SD media contained 2% agar (Bacto-Agar; Fisher). Standard yeast transformation, gene transplacement and other genetic methods have been previously described.(Ito et al., 1983; Rothstein, 1983) All site-directed mutagenesis experiments were performed using the New England Biolabs Q5 system according to manufacturer’s instructions. The identities of all constructs were confirmed by DNA sequencing (Eton Bioscience).

### Fatal attraction *sec14Δ* rescue assays

The viability of two haploid *sec14Δ* yeast strains of opposite mating type (MAT<u>a</u> vs MAT<u>α</u>) is supported by recessive loss-of-function *cki1^0^* and *kes1^0^* ‘bypass Sec14’ alleles, respectively.(Cleves *et al*., 1991; Fang et al., 1996) One of the pair is reconstituted for expression of a Sec14 variant of interest (Sec14*). Mating yields diploid cells where complementation of both ‘bypass Sec14’ alleles reimposes the essential Sec14 requirement for cell viability. Thus, only a functional Sec14 variant supports diploid cell viability. The appropriate *SEC14*, *SFH1* or *SFH1^E126A^*expression cassettes were linearized and recombined into the *LEU2* locus of strain CTY303 via transformation of yeast by the lithium acetate method and selection for histidine prototrophy.(Ito *et al*., 1983) Unselected Leu^−^ transformants were isolated, grown to an OD_600_ of ∼1.0 in YPD medium and mixed in a 1:1 ratio with strain CTY335 cultured similarly. Mixed cultures (1 mL total volume) were incubated without shaking overnight at 25°C. A 5 µL aliquot of each culture suspension was subsequently painted onto solid YPD medium, grown overnight, and the patches replica plated onto minimal glucose SD medium without additional supplements to select for diploid cells. Plates were inspected for viable diploids after 48 hours of incubation at 25°C. Recovery of viable diploid cells identified Sec14 mutants capable of functioning as a sole source of essential Sec14 activity in cells.

### Protein expression and purification

The His_8_-SUMO-Sec14 expression cassette was custom synthesized in pET28a+ (Genscript) and expressed as an N-terminal His_8_-SUMO fusion protein. *SFH1* open reading frame DNAs were subcloned into pET28b+ vectors as *Nco*I and *Sac*I restriction fragments such that an 8×histidine tag was incorporated at the N-terminus of each open reading frame. Expression constructs for Sec14^F219W^, Sec14^F221W^, Sec14^F228W^, Sfh1^F221W^, Sfh1^F223W^, Sfh1^V230W^, Sfh1^E126A^, and Sfh1^F223W,K197C,F233C^ were generated using the New England Biolabs Q5 system and confirmed by DNA sequencing. All proteins were produced in and purified from phage resistant *E. coli* BL21(DE3) cells. For ^19^F labeling, cells were grown using a resuspension method where the cultures were transferred from LB medium to minimal M9 medium supplemented with 60 mg of 5-fluoro-L-tryptophan prior to induction of protein expression.

Sfh1 proteins were expressed and purified as previously described.(Chen *et al*., 2023) His_8_-SUMO-Sec14 protein expression was induced by 0.5 mM IPTG and carried out overnight at 16 °C. The cells were harvested and lysed by passage through a French press in a buffer containing 25 mM Tris at pH 7.5, 500 mM NaCl, appropriate dilution of the HALT EDTA-free protease inhibitor cocktail, 0.2 mg/mL lysozyme, and a flake of DNase. His_8_-SUMO-Sec14 proteins were occupied with PtdCho in clarified cell lysates via addition of 25 mg of sonicated POPC per 50 mL of lysate and incubation at 30 °C for 1 hour. Fusion proteins were purified by Ni-NTA affinity chromatography and subjected to SUMO cleavage as previously described.(Katti et al., 2022) A second Ni-NTA affinity purification step separated Sec14 from the His-tagged SUMO and SUMO protease. All Sec14 and Sfh1 proteins were filtered through a 16/60 Sephacryl S-100 column in 25 mM sodium phosphate at pH 7.5, 300 mM NaCl, 5 mM β-mercaptoethanol (2ME), and 1 mM NaN_3_ as final polishing step. Fractions corresponding to monomeric proteins were pooled, concentrated and used for subsequent experiments.

### Crystallization and structure determination of the Sec14::PtdCho, Sec14::PtdCho::NPPM481, and Sfh1^F223W,K197C,F233C^::PtdEtn complexes

All proteins were crystallized using a hanging-drop vapor diffusion method at 16 °C (Sec14 complexes) or 23 °C (Sfh1^F223W,K197C,F233C^::PtdEtn). Purified binary Sec14::PtdCho complex was buffer exchanged into 10 mM HEPES at pH 7.2, 100 mM KCl, 5 mM 2ME and concentrated to 850 µM. Sec14::PtdCho was crystallized from a precipitant containing 0.1 M Bis-Tris at pH 6.5 and 40% v/v PEG 400. The ternary Sec14::PtdCho::NPPM481 complex was prepared by adding two-fold molar excess of NPPM481 to a dilute Sec14::PtdCho solution followed by concentrating the mixture for crystallization trials. Sec14::PtdCho::NPPM481 was crystallized from a precipitant containing 0.05 M Bis-Tris at pH 6.5, 0.05 M ammonium sulfate and 30% v/v pentaerythritol ethoxylate. Purified binary Sfh1^F223W,K197C,F233C^::PtdEtn complex was buffer exchanged into 20 mM HEPES at pH 7.2, 300 mM KCl and concentrated to 250 µM. Sfh1^F223W,K197C,F233C^::PtdEtn was crystallized from a precipitant containing 0.1 M Bis-Tris at pH 5.5 and 0.1 M ammonium sulfate.

The cryoprotectants used for crystal flash freezing in liquid nitrogen were as follows: 25% (v/v) glycerol in mother liquor for Sec14:PtdCho:NPPM481; 20% (v/v) glycerol in 0.05 M Bis-Tris at pH 5.5, 20% v/v PEG 400 for Sec14::PtdCho; and 30% (v/v) glycerol in 0.07 M Bis-Tris at pH 5.5, 0.07 M ammonium sulfate for Sfh1^F223W,K197C,F233C^::PtdEtn.

Diffraction data were collected at the U.S. Department of Energy’s (DOE) Brookhaven National Laboratory, New York using NSLS-II Beamline 17-ID-1. Data were indexed, integrated and scaled using the XDS software package.(Kabsch, 2010) The structures were solved by molecular replacement with PHASER(McCoy et al., 2007) and MOLREP(Vagin and Teplyakov, 2010) from the CCP4i suite.(Agirre et al., 2023; Potterton et al., 2003) The search models were as follows: Sfh1::PtdCho (PDB 3B7Q) for Sec14::PtdCho, Sfh1::PtdEtn (PDB 3B74) for Sfh1^F223W,K197C,F233C^::PtdEtn and Sec14::picolinamide (PDB: 6F0E) for Sec14::PtdCho::NPPM481. Manual model building and refinement were carried out with COOT(Emsley and Cowtan, 2004) and PHENIX 1.20.1(Afonine et al., 2012), respectively. Ligand restraints were generated with ELBOW.(Moriarty et al., 2009)

The quality of the final structural models was assessed using the wwPDB Validation Server. Visual Molecular Dynamics (VMD),(Humphrey et al., 1996), UCSF Chimera,(Pettersen et al., 2004) CCG Molecular Operating Environment (MOE) (Chem. Comp. Group Inc., Montreal, Canada), and LigPlot^+^(Laskowski and Swindells, 2011) were used for structural analyses and figure preparation. The coordinates and structure factors were deposited in the Protein Data Bank with accession codes 9YV3 for Sec14::NPPM481::PtdCho, 9EFP for Sec14::PtdCho, and 9EFN for Sfh1^F223W,K197C,F233C^::PtdEtn.

### PtdIns-transfer assays

Transfer of [^3^H]-PtdIns between membrane bilayer systems was monitored using rat liver microsomes and PtdCho liposomes as donor and acceptor membranes, respectively, and purified proteins as previously described.(Schaaf *et al*., 2008) All PtdIns-transfer assays were incubated for 30 min at 37°C. For the data in Figure 2F, input [^3^H]-PtdIns ranged from 14952-16648 cpm, experimental backgrounds from 216-294 cpm, and transfer for wild-type Sec14 was 12.2-15.4%. For the PtdIns-transfer data in Figure 4B, input [^3^H]-PtdIns ranged from 10,043 - 11,872 cpm, experimental backgrounds from 204-289 cpm, and transfer for wild-type Sfh1 was 9.1-13.1%. The data shown in Figure 7E represent the averages and standard deviations from 3 independent experiments and compared to wild-type transfer values measured for 140 pmoles Sec14 set at 100%. Input [^3^H]-PtdIns ranged from 9,135 - 11,118 cpm, experimental backgrounds ranged from 707-988 cpm, and transfer ranged from 10.2 to 17.8%.

### Hydrogen deuterium exchange coupled with mass spectrometry

Samples were prepared from 250 µM stock solutions of PtdCho-associated Sec14/Sfh1/Sfh1^E126A^ with and without 116 µM for DMPC/DDMB bicelles (30 mM total lipid). Stock solutions were equilibrated at 30°C for 20 min before a 1:9 dilution with H_2_O or D_2_O buffer. The final buffer was 10 mM HEPES at pH/pD 7.2, 300 mM KCl, and 2 mM TCEP. Exchange proceeded at 30°C for 10, 10^2^, 10^3^, 10^4,^ and 10^5^ s, and was quenched by mixing 3 μL of sample with 97 μL of a cooled mix of buffer and quench (0.94 parts buffer and 1 part quench of 0.8% (v/v) formic acid, 1.6 M guanidinium) before flash freezing in liquid nitrogen. Samples were stored at −80 °C.

Samples were thawed immediately before injection into a Waters™ HDX manager in line with a SYNAPT G2-*Si*. Samples were digested by *Sus scrofa* pepsin A (Waters™ Enzymate BEH) at 15°C and subsequently injected onto an online size exclusion column for lipid removal (Waters^TM^ Acquity UPLC Protein BEH SEC) and trapped on a C4 pre-column (Waters™ Acquity UPLC Protein BEH C4) at 1°C. The flow rate was 100 μL/min for 3.4 min. The chromatography buffer was 0.1% (v/v) formic acid. Peptides were subsequently resolved by binding to a C18 column (Waters™ Acquity UPLC BEH) at 1°C and eluted with a linear 3-40% (v/v) acetonitrile gradient using a flow rate of 40 μl/min for 7 min. Samples were injected in a random order. Mass spectrometry data were acquired using positive ion mode, HDMS^E^, and resolution mode. Peptide precursor and fragment data were collected via collision-induced dissociation at low (6 V) and high (ramping 22-44 V) energy. Capillary voltage was set to 2.4 kV for the sample sprayer. Desolvation gas was set to 650 L/h at 175°C. The source temperature was set to 80°C. Cone and nebulizer gases were flowed at 90 L/h and 6.5 bar, respectively. The sampling cone and source offsets were both set to 30 V. Data were acquired at a scan time of 0.4 s with a range of 100–2000 m/z. Mass correction used [Glu1]-fibrinopeptide B as a reference. For ion mobility, the wave velocity was 700 ms^−1^ and the wave height was 40 V.

Raw data from H_2_O-only controls were processed by PLGS (Waters™ Protein Lynx Global Server 3.0.3) using a database containing *S. scrofa* pepsin A, Sec14, Sfh1, and Sfh1^E126A^. In PLGS, the minimum fragment ion matches per peptide were 3, and methionine oxidation was allowed. The low and elevated energy thresholds were 250 and 50 counts, respectively, and the overall intensity threshold was 750 counts. DynamX 3.0 was used to search deuterated samples for peptides with 0.3 products per amino acid and one consecutive product. Structural images were made using residue-level scripts from DynamX in PyMOL without statistical filtering (Schrödinger, http://www.pymol.org/pymol), and heatmaps were created using in-house Python scripts with a minimum difference and significance filter (*p*-value <0.01 in a Welch’s t-test) applied.(Zhang et al., 2021) All samples had >3 replicates, except for Sec14 alone exchanged for 100 s (two replicates).

Data statistics and peptide lists are summarized in **Table S2**. The theoretical maximum deuteration used in percent calculations was determined as follows: (residues in peptide – 1 for N-terminal residue – the number of prolines not at the N-terminal position). Back exchange was calculated using peptides from samples that had plateaued (<2% difference in deuteration at 10^4^ and 10^5^ s) and had >40% deuteration. The mass spectrometry proteomics data are deposited with the ProteomeXchange Consortium via the PRIDE(Perez-Riverol et al., 2025) partner repository with the dataset identifier PXD066325.

### Sample preparation for ^19^F NMR spectroscopy

All protein samples for ^19^F NMR experiments were prepared in NMR buffer containing 150 mM NaCl, 150 mM arginine, 150 mM glutamic acid, 10 mM MES at pH 6.3, 0.02% NaN_3_, 8% D_2_O, and 1 mM TCEP. TCEP was omitted from buffer in samples for PRE experiments. Protein concentrations in NMR samples ranged from 45 to 150 µM.

Isotropically tumbling DMPC/DHPC bicelles (q=0.5) were prepared as described.(Katti, 2020) In brief, aliquots of DMPC and DHPC solutions in chloroform were dried under vacuum. The lipid films were hydrated and resuspended in NMR buffer to achieve a 1:2 DMPC:DHPC molar ratio. Three freeze-thaw cycles were performed to generate a clear bicelle stock solution containing 100 mM DMPC and 200 mM DHPC. Total lipid concentration was quantified by assay for phosphate.(King, 1932). The final total lipid concentration in all NMR samples was 80 mM, or *ca.* 310 µM of bicelle particles. For PRE experiments, 620 µM of 5-doxyl-PtdCho were dried and resuspended with mixtures of Sfh1/Sec14 variants and 80 mM total lipid.

### 1D ^19^F NMR experiments

All NMR experiments were conducted on an Avance III Neo NMR instrument (Bruker BioSpin, Billerica, MA) operating at the ^1^H Larmor frequency of 600 MHz and equipped with a Prodigy cryoprobe. The temperature was calibrated using methanol-d_4_ and set at 25°C. 1D ^19^F NMR spectra were collected using a spectral width of 20 ppm and an ^19^F carrier frequency at - 123.0 ppm. The recycle delay was set to 4 s in all experiments.

NMR spectra of bicelle-free and bicelle-containing samples were collected with 256-4096 and 4096-8192 scans, respectively. All data processing and analyses were performed using the MestReNova software package, v.14.2.0. The ^19^F chemical shifts were referenced to the external standard α,α,α -trifluorotoluene (Sigma-Aldrich) that resonates at −63.72 ppm relative to CFCl_3_ at 0 ppm. A sharp peak near −120 ppm observed in buffer-only samples corresponded to free fluoride leaching from the glass NMR tubes. This artifact was eliminated by subtracting the buffer-only spectra from corresponding ^19^F NMR spectra of protein-containing samples. The FIDs were apodized with the 50 or 70 Hz Gaussian function and zero-filled twice prior to the Fourier transform.

### 2D ^19^F-^19^F exchange spectroscopy (EXSY)

2D ^19^F-^19^F exchange spectroscopy (EXSY) experiments were carried out at 35°C. The temperature was calibrated using ethylene glycol. NMR samples contained 150 µM Sfh1 variants and DMPC/DHPC bicelles with a total lipid concentration of 80 mM. 2D ^19^F spectra were collected with mixing times t_mix_ of 4 µs, 20 ms, 50 ms, 100 ms, 150 ms, 250 ms, and 350 ms. The data were processed with NMRPipe(Delaglio et al., 1995) and analyzed with Sparky.(Lee et al., 2015) The interconversion between the two conformational states a and b of ^19^F-W_223_ or ^19^F-W_230_ manifested itself in the time-dependent buildup of the ^19^F cross-peaks. The intensities of the auto-peak I_aa_ and the cross-peak I_ba_ in each 2D spectrum were determined, and their ratio was plotted as a function of mixing time. The data were fit using the following equation(Pantoja et al., 2022) to obtain the forward and reverse rate constants, k_1_ and k_-1_:

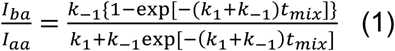

### 1D ^19^F relaxation dispersion experiments

The pulse sequence for constant-time ^19^F-detected CPMG relaxation dispersion experiments was adapted from previously published work.(Overbeck et al., 2020) Relaxation experiments were conducted at 35 °C on the F_223_W and V_230_W Sfh1 mutants with ^19^F carrier frequencies placed at −125.5 ppm and −125.6 ppm to capture the dispersion behaviors of the ^19^F-W_223_^Sfh1^ and ^19^F-W_230_^Sfh1^ peaks, respectively. The spectral width was set to 20.1 ppm, and the number of scans ranged from 2048 to 4096 scans per CPMG frequency point. The ^19^F-W_223_^Sfh1^ data were collected with a relaxation delay T_CPMG_=2 ms and 10 CPMG frequencies with 3 duplicates (500, 500, 1000, 1500, 2000, 2500, 2500, 3000, 3500, 4000, 4500, 5000, 5000 Hz). The ^19^F-W_230_^Sfh1^ data were collected with a relaxation delay T_CPMG_=3 ms and 12 CPMG frequencies with 3 duplicates (333.3, 333.3, 666.7, 1000.0, 1333.3, 1666.7, 1666.7, 2000.0, 2333.3, 2666.7, 3333.3, 3666.7, 4333.3, 5000.0, 5000.0 Hz).

The data were apodized with a 100 Hz Gaussian function, zero-filled to 2048 points and Fourier-transformed using the MestReNova software package (v.14.2.0). The effective transverse relaxation rates R_2,eff_ were calculated as:

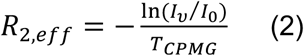

where I_ν_ is the intensity of the ^19^F resonance at CPMG frequency ν (ν_CPMG_), and I_0_ is the intensity of the reference spectrum obtained without the relaxation period. Uncertainties in R_2,eff_ values were estimated from the 3 duplicate ν_CPMG_ measurements. CPMG dispersion profiles were constructed by plotting R_2,eff_ values as a function of ν_CPMG_. Dispersion curves were fit with a two-state exchange model described by Equation 25 in Loria et al.(Palmer III *et al*., 2001) The following parameters were extracted from the fit: free-of-exchange transverse relaxation rate R_2,0_, exchange rate k_ex_, the chemical shift difference between the conformers Δω, and populations of the exchanging species p_A_ and p_B_. Uncertainties in the extracted parameters are given by the standard deviations over 100 fitting iterations to datasets generated via Monte Carlo simulations in which the data points are varied according to a Gaussian distribution. The width of this Gaussian noise was determined from the duplicate data points measured at three CPMG frequencies. Custom MATLAB scripts were developed and used for data analysis.

### Molecular dynamics (MD) simulations

The crystal structures of Sec14::PtdCho (PDB ID 9EFP) and Sfh1::PtdCho (PDB ID 3B7Q) were used as starting protein–lipid complexes. Four independent 2 µs replica simulations (denoted as r1 through r4) were performed for each complex, with the protein–lipid complex initially placed in solution approximately 90 Å above the bilayer center in four different orientations. Systems containing either Sec14::PtdCho or Sfh1::PtdCho, a lipid bilayer with 128 lipids per leaflet (90 POPC, 25 POPI, and 13 POPE), and 150 mM K⁺ and Cl⁻ ions were assembled using the CHARMM-GUI Membrane Builder.(Wu et al., 2014) Topologies and parameters for the phospholipid ligands were generated with the CGenFF server,(Vanommeslaeghe and MacKerell Jr, 2012) and the systems were solvated with TIP3P water(Jorgensen et al., 1983) in a rectangular simulation box. The CHARMM36m force field(Huang et al., 2017) was used in CHARMM GUI to generate coordinate, topology and parameter files. All system details are provided in **Table S3**.

MD simulations were carried out with GROMACS 2023.(Abraham et al., 2015) Nonbonded interactions were computed using the Verlet neighbor-searching scheme with a real-space cut-off of 1.2 nm. Van der Waals interactions were truncated at 1.2 nm, and electrostatic interactions were treated with the Particle Mesh Ewald (PME)(Darden et al., 1993) method using the same cut-off. Covalent bonds were restrained with the LINCS algorithm.(Hess et al., 1997) Protein-membrane systems were equilibrated using the default seven-step protocol that consists of the energy-minimization step, two NVT simulations, and four NPT simulations. Energy minimization employed the steepest-descent algorithm with strong positional restraints on the protein and lipid headgroups and dihedral restraints on selected torsion angles. The two subsequent NVT equilibration steps, 125 ps each, were performed with the V-rescale thermostat(Bussi et al., 2007) and relaxed positional restraints on the protein and lipids. The system was then equilibrated in four NPT ensembles (125, 500, 500, and 500 ps) using the V-rescale thermostat and C-rescale barostat.(Bernetti and Bussi, 2020) Semi-isotropic pressure coupling was applied to allow independent relaxation of box dimensions and membrane area. The positional and non-essential dihedral restraints on protein and lipids were gradually released during the four NPT steps to obtain a fully relaxed system for production runs.

MD production runs were performed at 310.15 K using the Nosé-Hoover thermostat(Hoover, 1985; Nosé, 1984) and the Parrinello-Rahman barostat.(Parrinello and Rahman, 1981) The temperature coupling groups were defined as protein::ligand, lipids and solvent (water and ions). The integration timestep was 2 fs with snapshots of coordinates written every 100 ps. An additional four independent 2.5 μs replica simulations (denoted as r5 through r8) were conducted for membrane-bound Sfh1 with frame 18,615 of the Sfh1::PtdCho r2 as initial configuration. For each replica the randomized starting velocities were assigned from a Maxwell-Boltzmann distribution.

For membrane-free simulations we used the Sfh1::PtdIns complex structure obtained after lipid exchange in the membrane-containing system. The Sfh1::PtdIns conformation was taken from frame 904 of replica r5 and placed in an 80×80×80 Å³ solvent box containing 150 mM K⁺ and Cl⁻ ions using CHARMM-GUI Solution Builder.(Jo et al., 2008) Energy minimization was carried out with the steepest descent algorithm until the maximum force was below 1000 kJ mol⁻¹ nm⁻¹. The system was then equilibrated for 125 ps in the NVT ensemble followed by 500 ps in the NPT ensemble using the V-rescale thermostat and C-rescale barostat, respectively. One MD production run of 2.5 µs was performed in the NPT ensemble with the V-rescale thermostat and C-rescale barostat employing isotropic pressure coupling. The randomized starting velocities were assigned from a Maxwell-Boltzmann distribution.

The trajectories were processed using GROMACS gmx trjconv command-line tool and visualized using VMD.(Humphrey *et al*., 1996) How trajectories were analyzed and relevant parameterizations are summarized in **Table S4**. The dynamics of membrane-bound Sec14 and Sfh1 were analyzed using the CONAN software package.(Mercadante et al., 2018) Three-dimensional protein representations were prepared, and molecular graphics were generated using UCSF ChimeraX(Meng et al., 2023) and VMD. Graphs and plots were generated using gnuplot and Matplotlib.(Hunter, 2007)

### Circular dichroism measurements

Protein concentrations were adjusted to 0.1 mg/mL in 25 mM phosphate buffer at pH 7.5 and 100 mM NaCl. Far-UV CD spectra were collected at 25 °C on a JASCO J-1500 spectropolarimeter (JASCO, Inc., Easton, MD, USA) equipped with a KOOLANCE temperature controller (KOOLANCE Inc., Auburn, WA, USA) using 1 mm path-length quartz cuvettes (Hellma Macro-cuvette 110-QS). For each protein, three scans were recorded from 190 to 250 nm and averaged (1 nm step size; 4 s/nm integration time). A buffer-only spectrum was acquired under identical conditions and subtracted from each dataset. Secondary-structure contents were independently estimated from the CD spectra using the pattern-recognition/k-nearest-neighbors approach (K2D3 web server).(Louis-Jeune et al., 2012)

## ACKNOWLEDGEMENTS

This work was supported by grants NIH R35 GM131804 and BE-0017 from the Robert A. Welch Foundation to V.A.B., NIH RO1 GM108998 and A-2194-20240404 from the Robert A. Welch Foundation to T.I.I., R35GM133751 to S.D., and the Bankaitis Fund for Young Scientists. AK was supported by an Predoctoral Fellowship NIH F30 AI189067. The molecular dynamics simulations were conducted using high performance research computing resources provided by Texas A&M High Performance Research Computing (HPRC). We thank Profs. Inna Krieger and James Sacchettini (Texas A&M University) for their assistance in collecting diffraction data in the early stages of this work. All diffraction data reported herein were collected at NSLS-II Beamline 17-ID-1 at Brookhaven National Laboratory (New York, U.S.A), a U.S. Department of Energy (DOE) Office of Science User Facility operated for the DOE Office of Science by Brookhaven National Laboratory under Contract No. DE-SC0012704. The BNL Center for BioMolecular Structure (CBMS) is primarily supported by the National Institutes of Health, National Institute of General Medical Sciences (NIGMS) through a Center Core P30 Grant (P30GM133893), and by the DOE Office of Biological and Environmental Research (KP1605010).

## DECLARATION OF INTERESTS

The authors declare that they have no conflicts of interest. Current affiliations for the following are listed:

Xiao Ru Chen, Department of Physiology & Biophysics, Columbia University Irving Medical Center Savana Green, Department of Microbial Pathogenesis and Immunology, College of Medicine, Texas A&M Health Science Center

Danish Khan, Department of Biochemistry, Virginia Tech University

Joy Shaffer, Department of Microbiology, Univ. of Texas Southwestern Medical Center

Ashley Kidwell, Department of Microbial Pathogenesis and Immunology, College of Medicine, Texas A&M Health Science Center

## AUTHORS CONTRIBUTIONS

The work and the experimental strategies were conceived by V.A.B., T.I.I., and S.D’A. The manuscript was written by V.A.B., T.I.I. and S.D’A. with input from all authors. HDX experiments were designed by S.D’A, T.I.I. and V.A.B., conducted by M.C.C. and J.M.S., and data analyzed by M.C.C., J.M.S. and S.D’A. Fatal attraction assays, genetic complementation tests, protein stability assays and mutant construction were conducted by V.A.B., S.M.G., D.K., and P.I. In vitro PtdIns-transfer assays were performed by S.M.G., D.K. and A.K. X.C. and T.M. performed the NMR experiments and analyzed the data with T.I.I. The MD production runs were set up and executed by T.N., and analyses of the MD data were performed by T.N., T.I.I., and V.A.B. Crystallization conditions were identified and optimized by P.S. and S.M.G., I.V.K. collected the X-ray data, P.S. processed the data, and P.S. and S.M.G. built and refined the structures. Structural data were analyzed by P.S., S.M.G., T.I.I., V.A.B., I.V.K., and J.C.S. Circular dichroism analyses were performed and analyzed by K.Z. and P.S.

