## Supplementary Materials for "An Atomistic Description of Heterotypic Lipid Exchange by Sec14-like Phosphatidylinositol Transfer Proteins"

**FIGURES**

**Figure S1**

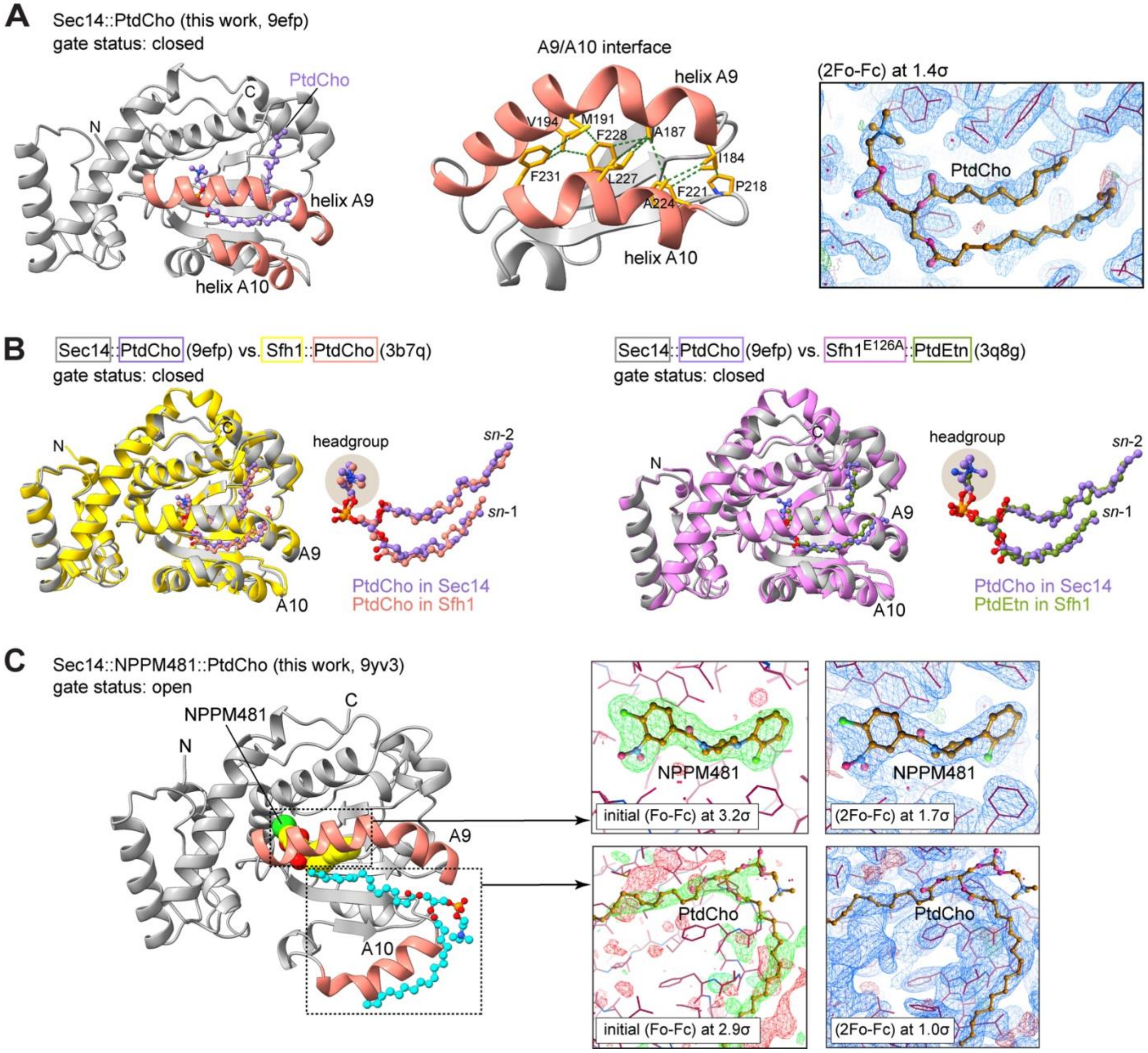

**Figure S1. Sec14::PtdCho and Sec14::NPPM481::PtdCho high-resolution structures and their analysis. (A)** High-resolution structure of the Sec14::PtdCho complex (gray), with helices A9 and A10 highlighted in salmon. Key Van der Waals A9/A10 interactions that stabilize the closed gate conformation are depicted with green dashed lines. Also shown is the (2Fo-Fc) map of bound PtdCho. **(B)** Backbone superposition of the Sec14::PtdCho structure (gray) with that of the Sfh1::PtdCho complex (left panel, yellow, r.m.s.d. 0.93 Å) and Sfh1$^{E126A}$::PtdEtn (right panel, pink, r.m.s.d. 0.95 Å). The poses of the bound lipid ligands are very similar in all binary protein::lipid complexes. **(C)** Structure of the Sec14::NPPM481::PtdCho ternary complex with NPPM481 and displaced PtdCho depicted in Van der Waals and ball-and-stick representations, respectively. (Fo-Fc) and (2Fo-Fc) electron density maps are shown for each ligand.

**Figure S2**

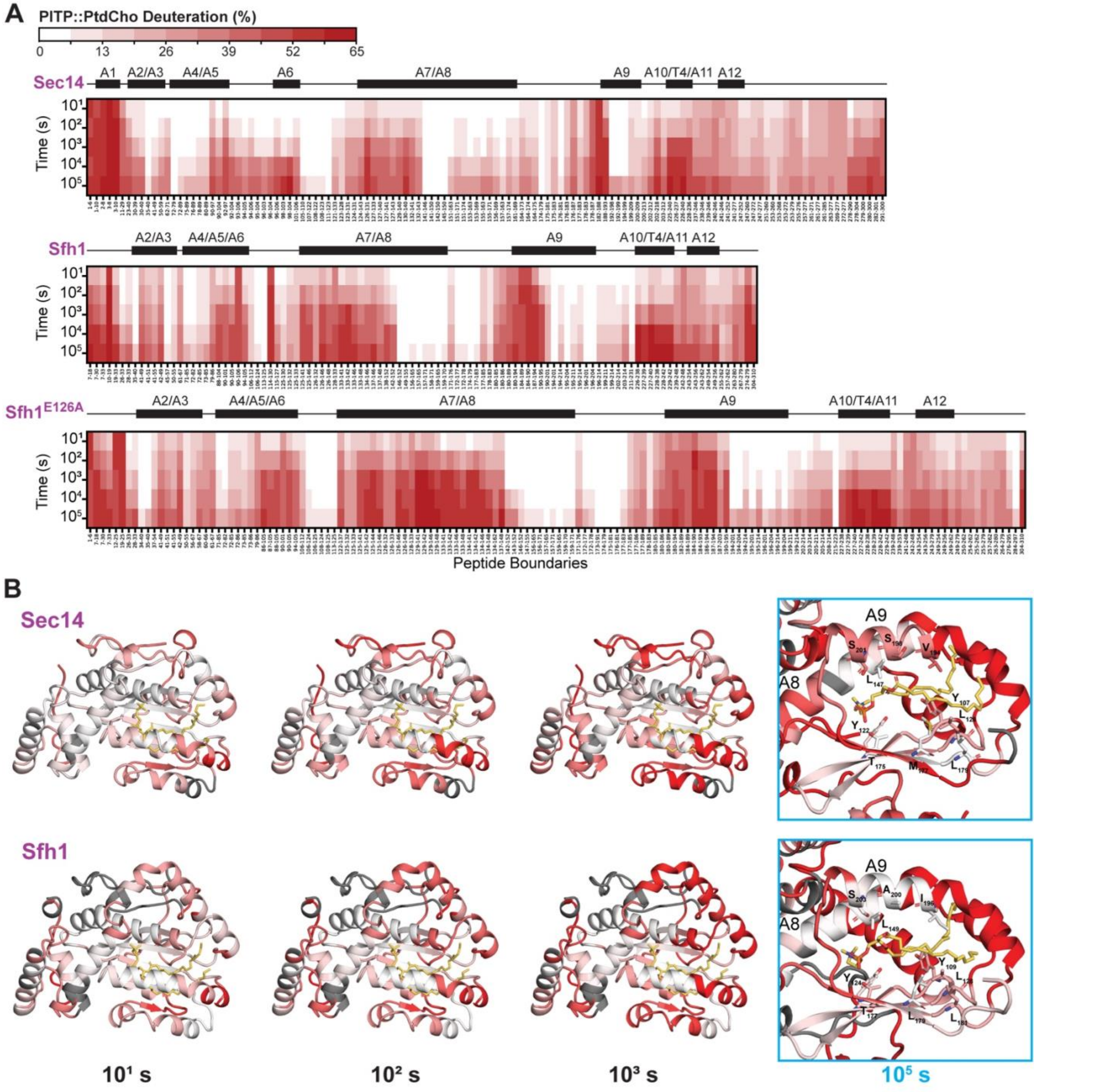

**Figure S2. Deuteration of PITP::PtdCho proteins in membrane-free systems. (A)** Deuteration of Sec14::PtdCho (top), Sfh1::PtdCho (middle), and $Sfh1^{E126A}$::PtdCho (bottom) in solution at $10^1$-$10^5$ s exchange. Helices are numbered as in Sec14. **(B)** Deuteration after $10^1$, $10^2$, or $10^3$ s of exchange projected onto the backbone structures of Sec14::PtdCho (9EFP, top) or Sfh1:: PtdCho (3B7Q, bottom). White-to-red coloration indicates 0-50% deuteration. Residues without coverage are shown in dark grey, and bound lipid is shown as yellow sticks. Residue-level data were obtained from DynamX. The boxed image shows a zoom of the lipid-binding pocket after $10^5$ s of exchange. Residues with low deuteration involved in lipid binding are shown as sticks and labeled. Helices $A_{10}$ and $A_{11}$ are hidden.

**Figure S3**

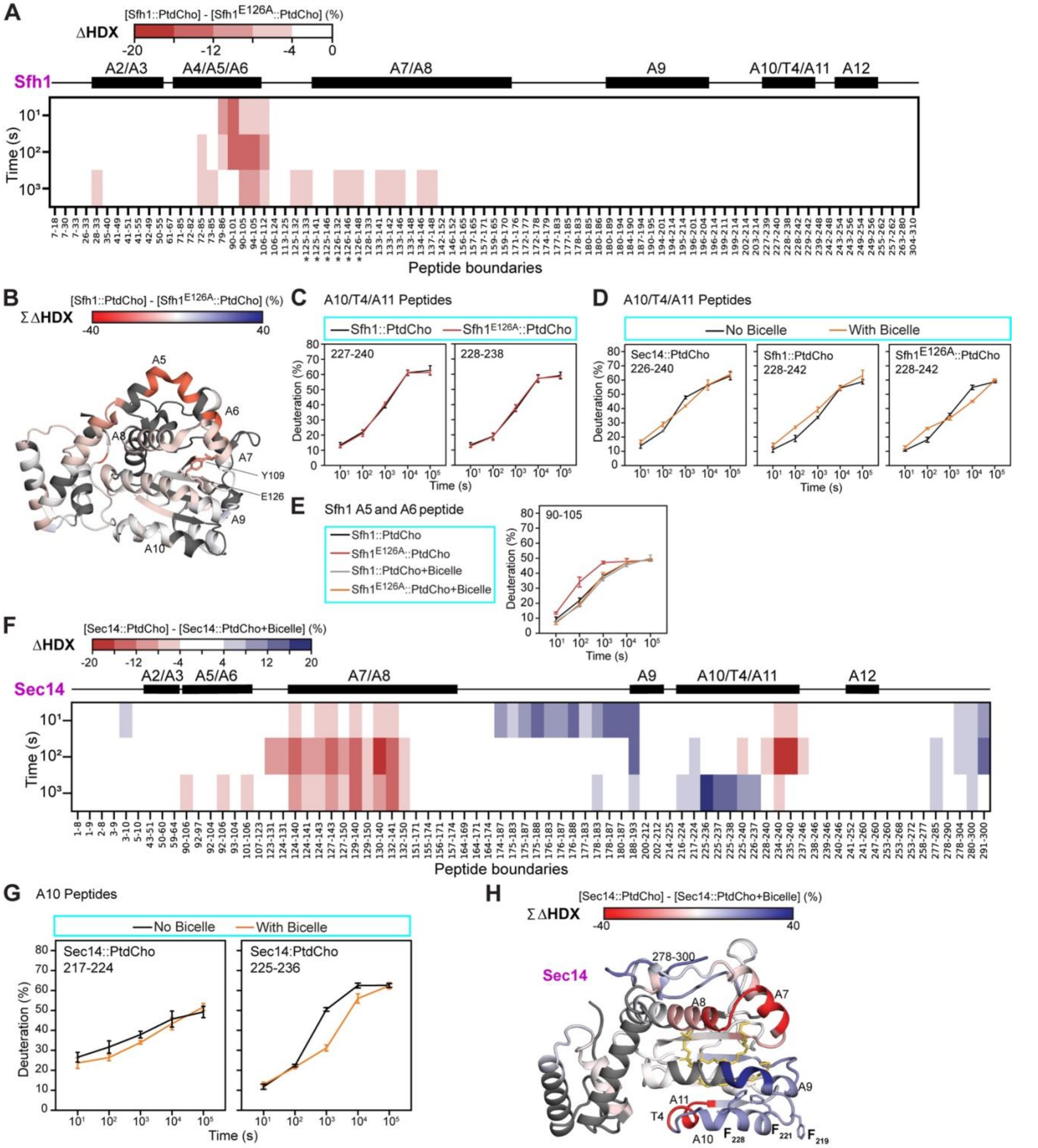

**Figure S3. Effect of the 'reactivating' $E_{126}$A mutation on Sfh1 conformational dynamics with and without bicelles. (A)** Changes in deuteration between Sfh1::PtdCho and $Sfh1^{E126A}$:: PtdCho. Peptides with ≥|4|% difference and a *p*-value ≤ 0.01 in a Welch's t-test (n=3) are indicated in red. Helices are numbered as in Sec14. Asterisks indicate peptides containing the $E_{126}$A mutation. **(B)** Increased deuteration as a result of the $E_{126}$A substitution is projected onto the Sfh1::PtdCho (3B7Q) structure. Changes are summed across the

10, $10^2$, and $10^3$ s time points. Residue-level data were obtained from DynamX with no statistical filtering. Residues without coverage are shown in dark grey. $E_{126}$ and $Y_{109}$ are shown as sticks. **(C-E)** Deuteration plots for representative peptides. Error bars are $\pm$ 2 S.D. (n=3 or 4), except for Sec14 alone at 100 s for which no error bars are shown as n=2 (grey cross). **(F)** Changes in deuteration of Sec14:: PtdCho upon the addition of bicelles (repeat of experiment in Fig. 2A, top panel). Peptides with ≥|4|% difference and a *p*-value ≤ 0.01 in a Welch's t-test (n=3) are indicated in red/blue. **(G)** Deuteration plots for representative $A_{10}$ peptides. Error bars are $\pm$ 2 S.D. (n=3). **(H)** Changes in deuteration upon addition of bicelles are projected onto the structure of Sec14:: PtdCho (9EFP). Changes were summed across the 10, $10^2$, and $10^3$ s time points. Residue-level data were obtained from DynamX with no statistical filtering. Residues without coverage are shown in dark grey. Bound lipid is in yellow and $F_{228}$, $F_{221}$ and $F_{219}$ are shown as sticks.

**Figure S4**

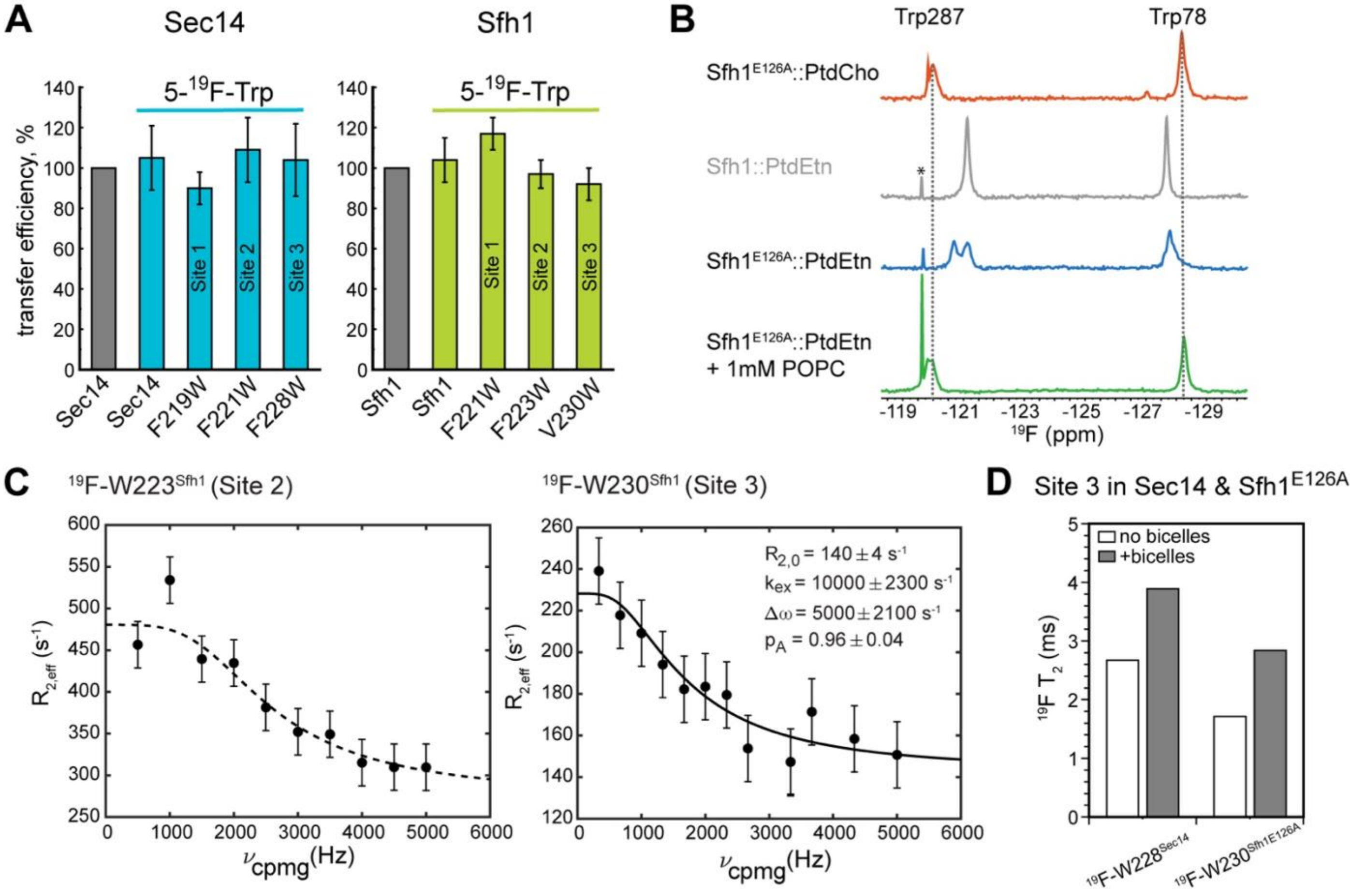

**Figure S4. Activity and NMR properties of 5-$^{19}$F-Trp labeled Sfh1, Sec14, and Sfh1$^{E126A}$ variants. (A)** PtdIns-transfer activities of the indicated purified recombinant Sec14 and Sfh1 proteins are shown. The specific variants are identified at bottom, and the $^{19}$F-Trp-labeled proteins are identified by the bar at top. Activities are related to those of unlabeled wild-type Sec14 and Sfh1 (each set at 100%; averages ± standard deviations, n=4). For Sec14 experiments, input [$^{3}$H]-PtdIns ranged from 19021-24699 cpm, background from 213-489 cpm, and transfer for wild-type Sec14 from 13.3-18.7%. For Sfh1 experiments, input [$^{3}$H]-PtdIns ranged from 7480-9200 cpm, backgrounds from 190-396 cpm, and transfer for wild-type Sfh1 from 8.1-10.6%. **(B)** Detection of PtdEtn exchange for PtdCho by Sfh1$^{E126A}$. The $^{19}$F spectra of 5-$^{19}$F-Trp labeled Sfh1$^{E126A}$ bound to PtdCho (red) and PtdEtn (blue). The chemical shifts of $W_{78}$ and $W_{278}$ $^{19}$F resonances depend on the chemical identity of the bound lipid ligand. Incubation of Sfh1$^{E126A}$::PtdEtn with POPC large unilamellar vesicles (LUVs, diameter 100 nm) switches the resonance signatures indicating lipid has been exchanged (green). Spectrum of the Sfh1::PtdEtn complex (gray) is shown as reference. Signal from trace fluorine contaminants in water is indicated (*). **(C)** $^{19}$F-based CPMG relaxation dispersion of $^{19}$F-$W_{223}^{Sfh1}$ (Site 2) and $^{19}$F-$W_{230}^{Sfh1}$ (Site 3) recorded in NMR buffer at 35 °C. Dashed line in the Site 2 plot is to guide the eye. Solid line in the Site 3 plot is the fit to the two-site exchange model (see Methods for details). **(D)** Transverse relaxation times, $T_2$, estimated from the linewidths of $^{19}$F-$W_{228}^{Sec14}$ and $^{19}$F-$W_{230}^{Sfh1E126A}$ in the absence (white) and presence of bicelles (gray). $T_2 = 1/(\pi \cdot \nu_{1/2})$, where $\nu_{1/2}$ is the linewidth at half-maximum in Hz.

**Figure S5**

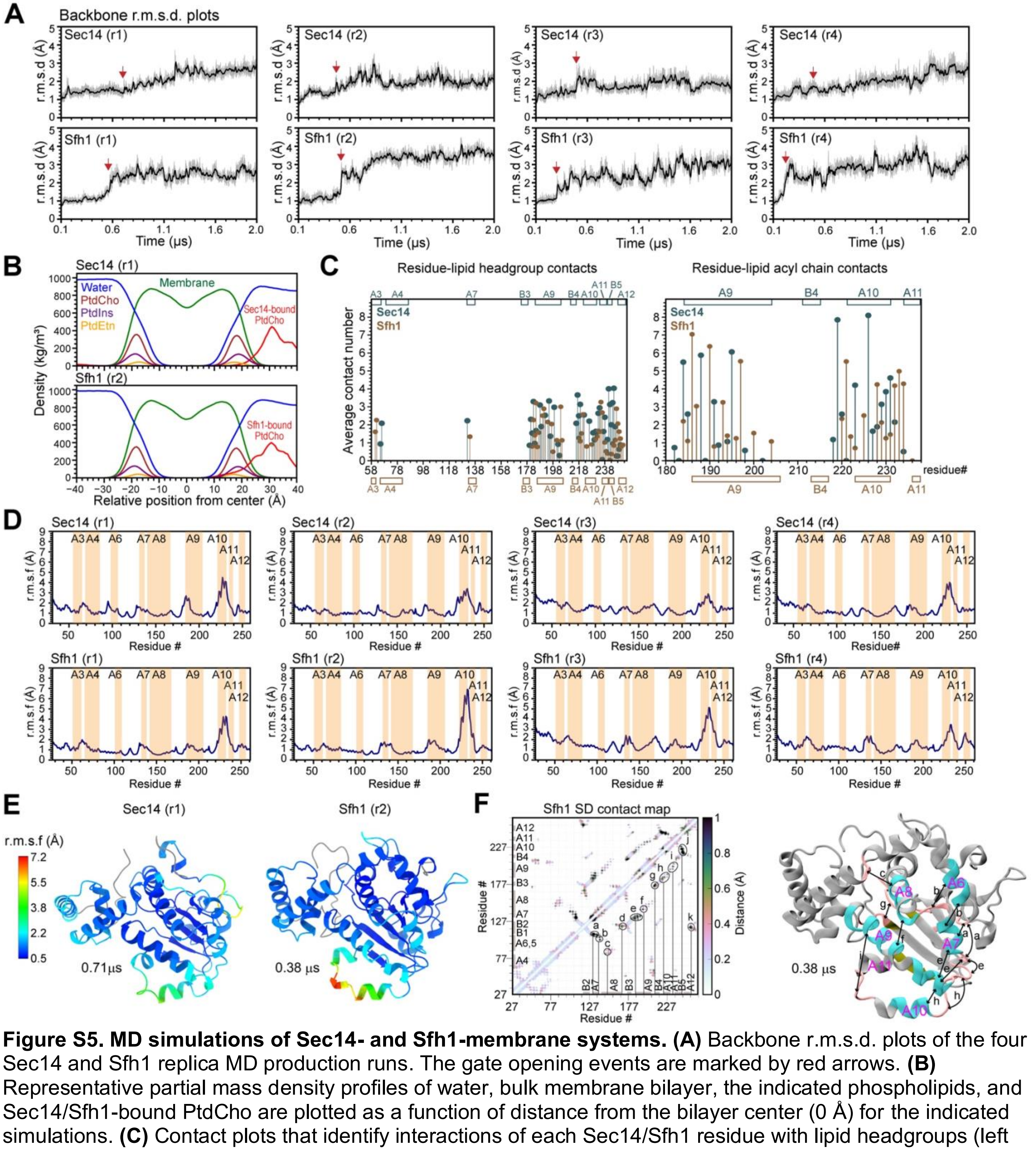

**Figure S5. MD simulations of Sec14- and Sfh1-membrane systems. (A)** Backbone r.m.s.d. plots of the four Sec14 and Sfh1 replica MD production runs. The gate opening events are marked by red arrows. **(B)** Representative partial mass density profiles of water, bulk membrane bilayer, the indicated phospholipids, and Sec14/Sfh1-bound PtdCho are plotted as a function of distance from the bilayer center (0 Å) for the indicated simulations. **(C)** Contact plots that identify interactions of each Sec14/Sfh1 residue with lipid headgroups (left panel) and acyl chains (right panel). Sec14 (slate gray) and Sfh1 (brown) secondary structure elements are identified at top and bottom, respectively. **(D)** Backbone r.m.s.f. plots of the four Sec14 and Sfh1 simulations. Selected helical regions are identified. **(E)** Backbone r.m.s.f. values are color-coded and mapped onto the MD snapshots of Sec14 and Sfh1. Frames used for the conformer representation are identified by simulation time and the replicas (r1 and r2). **(F)** Map of the r.m.s.d. values of inter-residue contacts in the membrane-bound Sfh1. Secondary structure elements (α-helices, cyan; β-strands, olive; loops, salmon) that experience relative pairwise fluctuations due to the motions of either one or both elements are marked by black arrows.

**Figure S6**

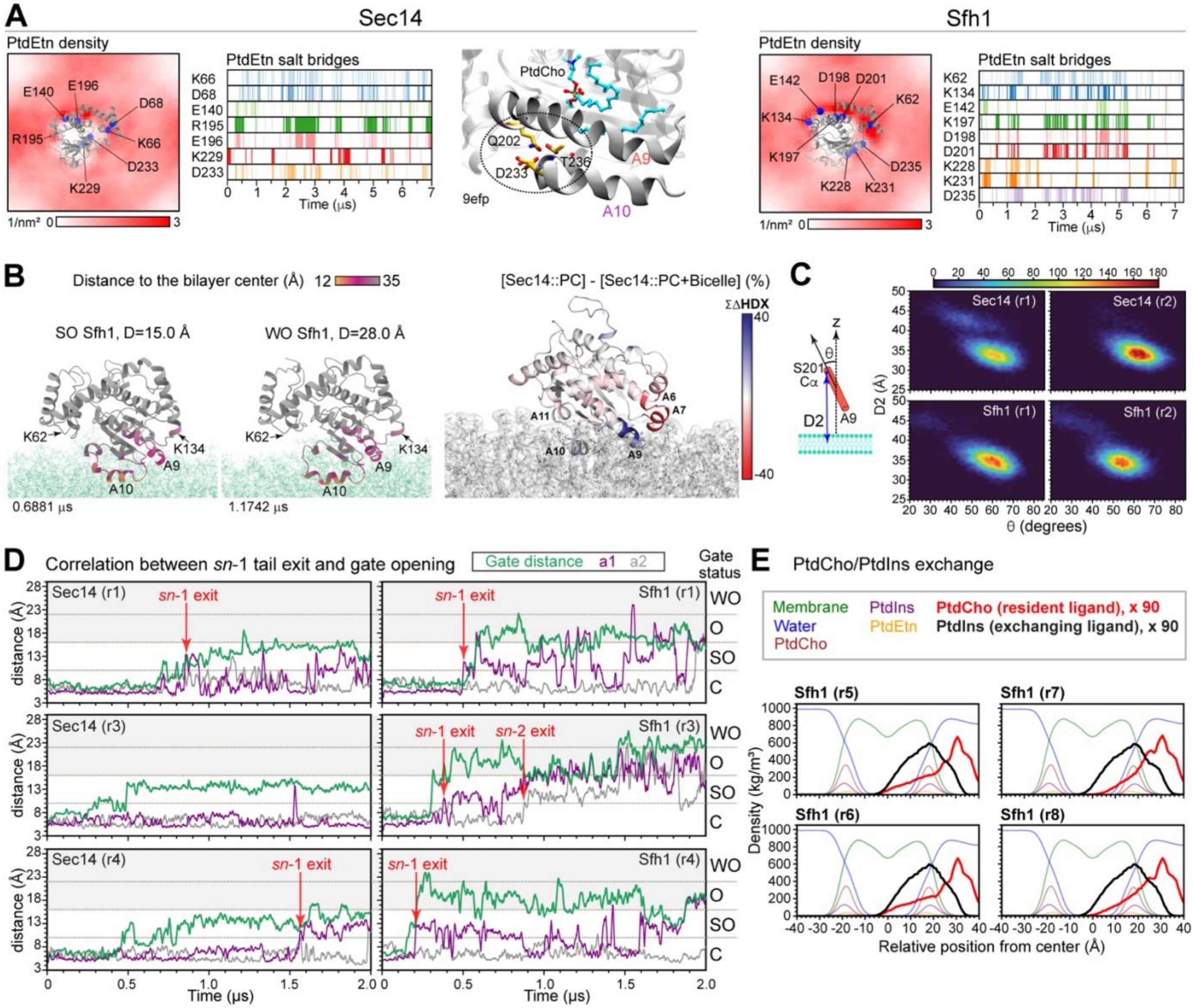

**Figure S6. PtdEtn demixing, protein geometry in membranes, and PtdCho exit. (A)** Demixing of PtdEtn by Sec14 (left panel) and Sfh1 (right panel) illustrated by two-dimensional lipid density plots and event plots for the most populated protein-PtdEtn salt bridges. Enlarged view of the closed Sec14 gate (middle panel) highlights polar residues $D_{233}$, $Q_{202}$ and $T_{238}$ (Sfh1 $D_{235}$, $Q_{204}$, $T_{238}$) whose interactions stabilize the closed gate conformation in the crystal structure. Residue $D_{233}$ interactions with PtdEtn might play a role in promoting gate opening. **(B)** Membrane-bound poses of semi-open (SO) and wide-open (WO) Sfh1 conformers with membrane insertion depth color-coded and mapped onto the MD snapshots from the Sfh1 r2 production run. The right panel shows the ΣΔHDX values (solution vs. bicelle-bound Sec14::PtdCho) mapped onto the MD snapshot of membrane-bound Sec14. **(C)** Protein geometry in membranes visualized using the distance D2 between Cα of Sec14 $S_{201}$/Sfh1 $S_{203}$ and bilayer center and the tilt angle θ of helix A9 relative to the membrane normal. Data are from the indicated replicas. **(D)** Gate ruler D, a1 and a2 distances plotted as a function of time for Sec14 and Sfh1 replica simulations. Points of PtdCho acyl chain *sn*-1 and *sn*-2 exit from the lipid binding pocket are indicated by arrows. Distances a1 and a2 are defined between the Cδ1 of Sec14 $L_{131}$/Sfh1 $L_{133}$ and methyl carbons of *sn*-1 and *sn*-2, respectively. **(E)** Partial mass densities calculated for replica simulations r5-r8 and for the indicated constituents are plotted as a function of distance to the bilayer center. The incoming

PtdIns and initially bound PtdCho partial mass densities are amplified 90 X in these plots. All replicas show exchange of PtdIns for PtdCho.

**Figure S7**

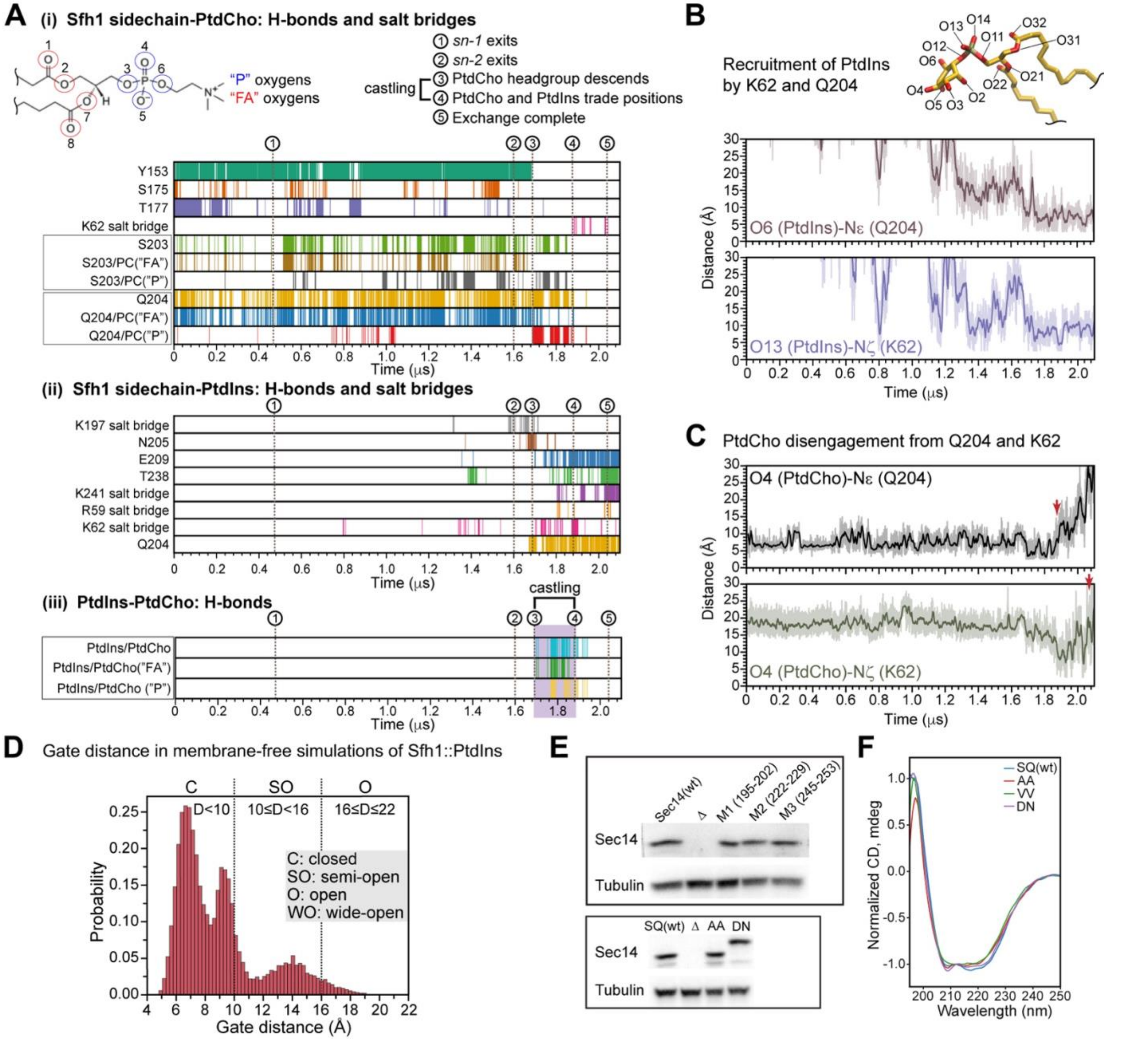

**Figure S7. Details of PtdCho/PtdIns exchange and properties of Sec14 mutants. (A)** Description of five key events of the PtdCho/PtdIns exchange reaction. The event plots highlight H-bond and salt bridge interactions that are formed/broken during exchange. The interacting partners are defined as (i) Sfh1-PtdCho, (ii) Sfh1-PtdIns, and (iii) PtdIns-PtdCho. **(B)** Recruitment of incoming PtdIns by interactions with residues $K_{62}$ of the RARKF motif and $Q_{204}$ is depicted as distance plots between the indicated amino acid and phospholipid groups. **(C)** PtdCho departure from the same residues highlighted in (B) is similarly plotted. Red arrows identify points of PtdCho disengagement from each. **(D)** Histogram of gate distance distributions in membrane-free simulations of the Sfh1:: PtdIns complex. **(E)** Immunoblot of Sec14 from yeast strains expressing the indicated variants (PtdIns clustering residue Ala-scan mutants at top; SQ motif mutants at bottom). The null condition was provided by the bypass Sec14 strain CTY303 deleted for the *SEC14* gene (see Methods). The $Sec14^{S201D,Q203N}$ mutant runs anomalously on SDS-PAGE. Tubulin immunoblotting served to normalize sample load. **(F)** Circular dichroism traces for Sec14 (SQ) and the indicated SQ mutant proteins (AA,DN,VV) are shown.

### TABLES

**Table S1.** Crystallographic data collection statistics and refinement values for Sec14::PtdCho, Sec14::NPPM481::PtdCho and Sfh1$^{K197C,F223W,F233C}$. Values in parentheses are for the highest resolution shell.

| **Data collection Parameters** | Sec14::PtdCho | Sec14::NPPM481::PtdCho | Sfh1$^{K197C,F223W,F233C}$ |
|---|---|---|---|
| Resolution range (Å) | 45.86- 1.83 | 30.71- 2.25 | 46.46- 1.75 |
| Space group | $P3_221$ | $P3_221$ | $P2_1$ |
| Unit cell parameters | | | |
| a, b, c(Å) | 74.4, 74.4, 195.8 | 89.2, 89.2, 100.4 | 49.4, 72.0, 114.6 |
| α, β, γ (°) | 90, 90, 120 | 90, 90, 120 | 90, 93.95, 90 |
| Number of molecules in the ASU | 2 | 1 | 2 |
| Vm (Å$^3$/Da) | 2.24 | 3.37 | 2.83 |
| Solvent Content (%) | 45.0 | 63.5 | 56.5 |
| Number of unique reflections | 56254 (5567) | 22389 (2189) | 80061 (7936) |
| Completeness (%) | 99.7 (99.9) | 100 (100) | 98.8 (98.6) |
| I/σ(I) | 5.9 (1.8) | 9.6 (2.1) | 7.2 (1.7) |
| $R_{meas}$ (%) | 14.3 (56.8) | 17.3 (187.3) | 14.2 (109.8) |
| $CC_{1/2}$ (%) | 98.5 (60.6) | 99.6 (82.9) | 99.3 (68.1) |
| **Refinement** | | | |
| $R_{cryst}$ (%) | 17.2 | 21.7 | 24.1 |
| $R_{free}$ (%) | 20.8 | 26.8 | 27.9 |
| R.m.s.d in bond lengths (Å) | 0.01 | 0.009 | 0.01 |
| R.m.s.d in bond angles (°) | 1.0 | 1.0 | 1.0 |
| R.m.s.d in torsion angles (°) | 16.8 | 14.4 | 12.2 |
| **Ramachandran plot analysis** | | | |
| Residues in the most favored regions (%) | 91.0 | 89.6 | 93.8 |
| Residues in the additionally allowed regions (%) | 8.6 | 10.0 | 5.9 |
| Residues in the generously allowed regions (%) | 0.4 | 0.4 | 0.4 |
| Residues in disallowed regions (%) | 0 | 0 | 0 |
| **Mean B factors (Å$^2$)** | | | |
| Wilson B factor | 28.9 | 48.7 | 20.7 |
| Protein atoms | 28.8 | 57.1 | 34.7 |
| Ligands | 35.4 | 52.6 | 51.1 |
| Solvent | 30.6 | 55.8 | 40.9 |
| Number of protein atoms | 4696 | 2430 | 5132 |
| Number of water oxygen atoms | 293 | 193 | 553 |
| PDB ID | 9EFP | 9YV3 | 9EFN |

**Table S2.** Summary of HDX experiments.

| | **HDX data for PtdCho-bound Sec14, Sfh1, and Sfh1-E126A, with and without bicelle** | **HDX data for PtdCho-bound Sec14 with and without bicelle (Sec14-Repeat)** |
|---|---|---|
| **Samples** | Sec14::PtdCho, Sfh1::PtdCho, and Sfh1E126A::PtdCho +/- bicelle | Sec14::PtdCho +/- bicelle |
| **Buffer** | 10 mM HEPES pD 7.2, 300 mM KCl, 2 mM TCEP, 30°C | 10 mM HEPES pD 7.2, 300 mM KCl, 2 mM TCEP, 0.55% DMSO, 30°C |
| **Time** | 10, 100, 1000, 10000, 100000 s | 10, 100, 1000, 10000, 100000 s |
| **Controls** | Unlabeled Sec14::PtdCho, Sfh1::PtdCho, and Sfh1E126A::PtdCho +/- bicelle | Unlabeled Sec14::PtdCho +/- bicelle |
| **Replicates** | 3 or 4 (except Sec14 alone at 100 s has only 2) | 3 |
| **Sec14** | | |
| # Peptides | 124 | 76 |
| Sequence Coverage | 97.40% | 78.90% |
| Average Peptide Length/Redundancy | 12.50 / 5.10 | 12.29 / 3.89 |
| Global Significance Threshold | 3.02%* | 3.56%* |
| Mean Back Exchange | 50.60% | 42.90% |
| **Sfh1** | | |
| # Peptides | 105 | |
| Sequence Coverage | 88.10% | |
| Average Peptide Length/Redundancy | 12.10 / 4.49 | |
| Global Significance Threshold | 3.81%* | |
| Mean Back Exchange | 48.70% | |
| **Sfh1-E126A** | | |
| # Peptides | 147 | |
| Sequence Coverage | 96.10% | |
| Average Peptide Length/Redundancy | 12.32 / 6.08 | |
| Global Significance Threshold | 3.53%* | |
| Mean Back Exchange | 43.80% | |
| *Calculated as the propagated standard error of the mean. Implemented significance threshold of >4% with a p-value <0.01 in Welch's t-test (n=4). | | |

**Table S3.** Summary of molecular dynamics simulations. For the membrane-containing simulations, the membrane composition was 90 POPC, 25 POPI, and 13 POPE lipid molecules per leaflet.

| System | Replica | Protein orientation | System size (# atoms) | Rectangular solvent box (x, y, z; Å) | Ions | Ion concen-tration (mM) | Protein-membrane COM distance (Å) | Production run (μs) |
|---|---|---|---|---|---|---|---|---|
| Sec14/ membrane | 1 | First principal axis and Z axis aligned | 145,079 | 92.7, 92.7, 178.9 | K+, Cl- | 150 | 90 | 2.0 |
| | 2 | +45° rotation about X | 142,040 | 92.7, 92.7, 175.5 | K+, Cl- | 150 | 90 | 2.0 |
| | 3 | −45° rotation about X | 137,166 | 92.7, 92.7, 170.4 | K+, Cl- | 150 | 90 | 2.0 |
| | 4 | +45° rotation about Y | 136,638 | 92.7, 92.7, 169.8 | K+, Cl- | 150 | 90 | 2.0 |
| Sfh1/ membrane | 1 | First principal axis and Z axis aligned | 142,089 | 92.7, 92.7, 176.1 | K+, Cl- | 150 | 89 | 2.0 |
| | 2 | +45° rotation about X | 144,163 | 92.7, 92.7, 177.7 | K+, Cl- | 150 | 92 | 2.0 |
| | 3 | −45° rotation about X | 143,243 | 92.7, 92.7, 177.2 | K+, Cl- | 150 | 92 | 2.0 |
| | 4 | +45° rotation about Y | 137,752 | 92.7, 92.7, 170.9 | K+, Cl- | 150 | 91 | 2.0 |
| | 5-8 | Frame 18,615 of replica 2 | 144,163 | 92.7, 92.7, 177.7 | K+, Cl- | 150 | 39 | 2.5 each |
| Sfh1::POPI in solution | 1 | Frame 904 of replica 5, membrane removed | 47,979 | 80.0, 80.0, 80.0 | K+, Cl- | 150 | N/A | 2.5 |

**Table S4**. MD trajectory parameterization and analysis.

| No. | Processing/Analysis | Atom Selection/Parameterization | GROMACS/ VMD tool |
|---|---|---|---|
| 1 | Removal of periodic boundary conditions | System | gmx trjconv (-pbc whole, -pbc nojump, -center, -ur compact, -pbc cluster) |
| 2 | Concatenation of trajectories | System | gmx trjcat (-settime) |
| 3 | Alignment of trajectory frames | Fitting group: Protein | gmx trjconv (-fit rot+trans) |
| 4 | Calculation of protein and ligand r.m.s.d. and r.m.s.f. | Protein backbone heavy atoms: N, Cα, and C;<br>Ligand heavy atoms | gmx rms<br>gmx rmsf |
| 5 | Helical gate distance D | Sec14: R195 Cα (A9) - F231 Cα (A10)<br>Sfh1: K197 Cα (A9) - F233 Cα (A10) | gmx distance |
| 6 | Protein-membrane Z distance | Sec14: F219 Cα and bilayer center<br>Sfh1: F221 Cα and bilayer center | gmx distance (-oxyz) |
| 7 | Partial mass density profiles | POPC::POPE::POPI, POPC, POPE, POPI, Water (TIP3), bound PtdCho, and incoming POPI (POPI505) | gmx density -center -sl 200 -d Z |
| 8 | Protein-resident PtdCho distances | Sec14:<br>L131 Cδ1 and C1 of resident PtdCho<br>L131 Cδ1 and C42 of resident PtdCho<br><br>Sfh1:<br>L133 Cδ1 and C1 of resident PtdCho<br>L133 Cδ1 and C42 of resident PtdCho<br>Q204 Nε and O4 of resident PtdCho<br>K62 Nζ and O4 of resident PtdCho | gmx distance |
| 9 | Sfh1-POPI505[a] distances | Q204 Nε and O6 of POPI505<br>K62 Nζ and O13 of POPI505 | gmx distance |
| 10 | Protein-lipid contacts | Distance cutoff: 6.0 Å<br>Contacts with lipid headgroups:<br>(Any atom of a protein residue) to (N of POPC, N of POPE, O4 of POPI).<br><br>Contacts with lipid acyl chains:<br>(Any atom of a protein residue) to (C23 of POPC, C23 of POPE, C23 of POPI).<br><br>Cutoff of the average number of contacts: 0.5 | gmx mindist |
| 11 | Helix A10 membrane insertion depth | Z distance between lipid bilayer center and:<br>F219 Cζ, F221 Cζ, F228 Cζ (Sec14)<br>F221 Cζ, F223 Cζ, V230 Cγ1 (Sfh1) | gmx distance (-oxyz) |

| No. | Processing/Analysis | Atom Selection/Parameterization | GROMACS/ VMD tool |
|---|---|---|---|
| 12 | Protein-lipid H-bonds | Donor-acceptor distance cutoff: 3.5 Å<br>Angle relaxation cutoff: 50°<br>Interacting partners: Sec14/Sfh1-PtdIns, Sfh1-PtdCho, PtdCho-POPI505, Sfh1-POPI505 | HBonds VMD plugin |
| 13 | Protein-lipid salt bridges | Interatomic distance cutoff: 3.5 Å<br>Interacting partners: Sec14/Sfh1-PtdIns, Sec14/Sfh1-PtdEtn, Sfh1-PtdCho, Sfh1-POPI505 | Salt Bridges VMD plugin |
| 14 | Membrane insertion depth of protein sidechains | Z distance between COM of protein side-chain heavy atoms and bilayer center | gmx distance (-oxyz) |
| 15 | 2D lipid density map (upper membrane leaflet) | PtdIns density calculations:<br>O2-O6, C11-C16, O11-O14, and P atoms of the POPI headgroup<br><br>PtdEtn density calculations:<br>N, C11-C12, O11-O14, and P atoms of the PtdEtn headgroup | gmx densmap -aver z -n1 50 -n2 50 |
| 16 | Protein-membrane encounter (distance vs. angle heatmap) | Z distance: between Cα of F219/F221 (Sec14/Sfh1) and lipid bilayer center<br><br>Angle: between the helix A9 vector and membrane normal. The A9 vectors are defined as Y188 Cα-R195 Cα in Sec14, and Y190 Cα-K197 Cα in Sfh1. | gmx distance (-oxyz)<br>gmx gangle -g1 vector -g2 z |
| 17 | Protein orientation in membrane (distance vs. angle heatmap) | Z distance: between Cα of S201/S203 (Sec14/Sfh1) and bilayer center<br><br>Angle: between the helix A9 vector and membrane normal. The A9 vectors are defined as Y188 Cα-R195 Cα in Sec14, and Y190 Cα-K197 Cα in Sfh1. | gmx distance (-oxyz)<br>gmx gangle -g1 vector -g2 z |

[a]POPI505 is the ID of the POPI molecule that replaces protein-bound Sec14.

### SUPPLEMENTARY VIDEOS

**Supplementary Video 1.** Stable membrane association of Sfh1 and gate opening. Key structural elements are highlighted; helices A6 and A7 in blue, A9 in orange and A10 and the preceding loop in purple. Residue $F_{221}$ is highlighted. PtdCho is in cyan.

**Supplementary Video 2.** The exchange of PtdIns for PtdCho by Sfh1. Close up view of the Sfh1 lipid binding pocket is shown from gate opening to castling and lipid exchange. Sfh1 backbone is in gray. Key amino acid side-chains are shown, bound PtdCho is in cyan ball & stick and helices A9 and A10 are highlighted in orange and purple, respectively. The RARKF backbone region is in lime green. Membrane lipids are not shown for purposes of clarity. Incoming PtdIns is in gold ball & stick.

**Supplementary Video 3.** Closure of the gate over newly incorporated PtdIns in a membrane-free system. Bound PtdIns is in gold ball & stick and helices A9 and A10 are highlighted in orange and purple, respectively. The RARKF backbone region is in lime green.
